# Hippocampal Glucuronyl C5-epimerase promotes stress resilience by directly engaging PI3K through a non-enzymatic mechanism

**DOI:** 10.64898/2026.05.02.722405

**Authors:** Meilin Chen, Lulin Huang, Songyu Yao, Jie Li, Fei He, Muya Xiong, Jianping Fang, Yanling Li, Yiwen Zhang, Wenfeng Liao, Zhenyun Du, Fei Guo, Tianyu Li, Jinjin Duan, Jing Nie, Yu Zhang, Can Jin, Yechun Xu, Yang Li, Xia Li, Yujun Wang, Jinggen Liu, Kan Ding

**Author notes:** These authors contributed equally to this work as co-first authors (Meilin Chen, Lulin Huang, Songyu Yao, Jie Li). Corresponding author (K.D.); (J.L.); (Y.W.); (X.L.).

## Abstract

Major depressive disorder (MDD) seriously affects human physical and mental health and causes global socioeconomic burdens. Stress-induced depressive etiology is linked to dendritic atrophy in the hippocampus, however, the underlying mechanism was poorly understood. Here, we identify that glucuronyl C5-epimerase (*Glce*), a heparan sulfae-modifying enzyme, as a critical regulator of hippocampal dendritic integrity and stress resilience. Genetic ablation of *Glce* in the hippocampal excitatory neurons is sufficient to induce dendritic atrophy and depressive-like behaviors, whereas its restoration rescues these deficits. Notably, *Glce* levels are reduced in the plasma of depressed individuals. We further find that *Glce* functions independently of its enzymatic activity, instead binding directly to and activating PI3K (p85α/p110α), thereby triggering an Akt/CREB/BDNF signaling cascade. Together, our findings uncover a non-canonical role for Glce and establish that *Glce*–PI3K–BDNF axis is essential for maintaining hippocampal structure and behavioral resilience, thereby highlighting a critical role of *Glce* in the pathophysiology of depression.

## Introduction

Major depressive disorder (MDD) is a debilitating psychiatric disorder characterized by the abnormal mood/emotion, cognition and behavior, seriously affecting human physical and mental health (Krishnan and Nestler, 2008; Malhi and Mann, 2018). A core pathophysiological feature observed in MDD patients is neuronal atrophy, particularly in the hippocampus, manifested as reduced volume and diminished synaptic plasticity (Ashtari et al., 1999). Preclinically, chronic stress, a major environmental risk factor for depression, similarly drives these structural deficits, leading to dendritic atrophy and synaptic loss in hippocampal neurons (Li et al., 2011; Magariños et al., 1996). The neurotrophic signaling, particularly through brain-derived neurotrophic factor (BDNF) is known to be crucial for maintaining dendritic integrity and promoting behavioral resilience (Duman and Voleti, 2012; Park and Poo, 2013; Yu et al., 2012). However, the detailed mechanisms underlying this maintenance and how they are disrupted by stress remain largely unknown.

Heparan sulfate proteoglycans (HSPGs), a family of cell surface and extracellular matrix proteins, are emerging as critical regulators of brain development and function (Condomitti and de Wit, 2018). Their biological activity is largely defined by the structural diversity of their heparan sulfate (HS) chains, which is dynamically shaped through enzymatic modifications including epimerization and sulfation (Blackhall et al., 2001; Esko and Selleck, 2002; Lindahl and Li, 2009). This enables HSPGs precise interactions with growth factors and morphogens to direct axon guidance, synaptogenesis, and plasticity (Inatani et al., 2003; Irie et al., 2012; Lee and Chien, 2004; Li et al., 2002; Zhang et al., 2018). Consequently, genetic disruption of HS-modifying enzymes impairs neural connectivity, as evidenced by axon guidance defects upon their conditional deletion and by synaptic deficits such as reduced spine density following loss of the enzyme SULF1 (Kalus et al., 2009). Notably, genetic variants in these enzymes and altered expression of HSPG core proteins are robustly linked to neurodevelopmental and psychiatric disorders (Condomitti and de Wit, 2018; Irie et al., 2012; Li et al., 2002). For example, mutations in the HS polymerase EXT1 are associated with susceptibility to autism spectrum disorder and schizophrenia (Li et al., 2002). These studies suggest the importance of HS modification in disease susceptibility. Intriguingly, a growing body of evidence suggests that some glycosaminoglycan-modifying enzymes may possess biological functions that are beyond their catalytic activity, serving as scaffolds or direct signaling modulators in specific cellular contexts (Cadwalader et al., 2012).

D-Glucuronyl C5-epimerase, encoded by *Glce* gene, is one of the key enzymes involved in the modification of HS (Li et al., 2001; Nadanaka and Kitagawa, 2008), responsible for conversion of D-glucuronic acid to L-iduronic acid (Hagner-McWhirter et al., 2004; Li et al., 2001), therefore influencing chain flexibility and binding capacity (Catlow et al., 2008; Jia et al., 2009; Ori et al., 2008). Studies with *C. elegans* demonstrate that mutation of alleles in the C5-epimerase gene results in the most severe defects on axonal development (Bülow and Hobert, 2004). Global deletion of *Glce* in mice results in severe embryonic or perinatal lethality (Feyerabend et al., 2006; Forsberg and Kjellén, 2001; Li et al., 2003; Perrimon and Bernfield, 2000). Despite this, the function of *Glce* in the adult brain, and specifically its potential involvement in stress-related depression has remained unknown.

In this study, we identified *Glce* as a novel and essential regulator of hippocampal dendritic morphology and resilience to depression. We demonstrate that *Glce* is required in hippocampal excitatory neurons to maintain dendritic integrity and to prevent the emergency of depression-like behaviors. Mechanistically, we discover that *Glce* exerts its protective effects not through its canonical enzymatic activity. Instead, Glce operates as a scaffold that directly binding to and activating the PI3K heterodimer (p85α/p110α), thereby initiating a signaling cascade through Akt/CREB to upregulate BDNF expression. In summary, our findings collectively reveal a non-canonical role for *Glce* and established a previous unrecognized *Glce*–PI3K–BDNF axis is essential for maintaining hippocampal structure and behavioral resilience. Furthermore, this study highlights *Glce* and its interaction with PI3K as promising therapeutic targets for future rapid-acting antidepressant therapies.

## Results

### Hippocampal *Glce* deletion induces depressive-like behaviors that are rescued by enzymatic-activity-independent *Glce* re-expression

Emerging evidence indicated the involvement of *Glce* in axonal development (Bülow and Hobert, 2004; Li et al., 2014) and neuropsychiatric disorders (Liachko et al., 2019), which lead us to explore its potential role in major depressive disorders (MDD). Given the critical role of the hippocampus in modulating stress response and mood (Anacker and Hen, 2017; Tartt et al., 2022), along with its high proportion of excitatory glutamatergic neurons (Yao et al., 2021), we first explored whether *Glce* is expressed in hippocampal glutamatergic neurons. We found that *Glce* is abundantly expressed in hippocampal excitatory glutamatergic neurons (Supplemental Figure 1, A and B), while minimal in GABAergic neurons or microglia (Supplemental Figure 1B).

To elucidate the function of *Glce* in glutamatergic neurons, a conditional knockout strategy was employed. By crossing a floxed *Glce* allele with a CaMKIIα Cre transgenic line, we generated mice with *Glce* deletion in targeted glutamatergic neurons, referred to as NKO mice (Figure 1A and Supplemental Figure 2, A to C). We found that these NKO mice displayed depressive-like behaviors, as evidenced by increased immobility in the tail suspension test (TST) and forced swim test (FST), and decreased sucrose consumption in the sucrose preference test (SPT), while their locomotion was not altered in the locomotor activity test (LAT) (Figure 1B).

**Figure 1.**
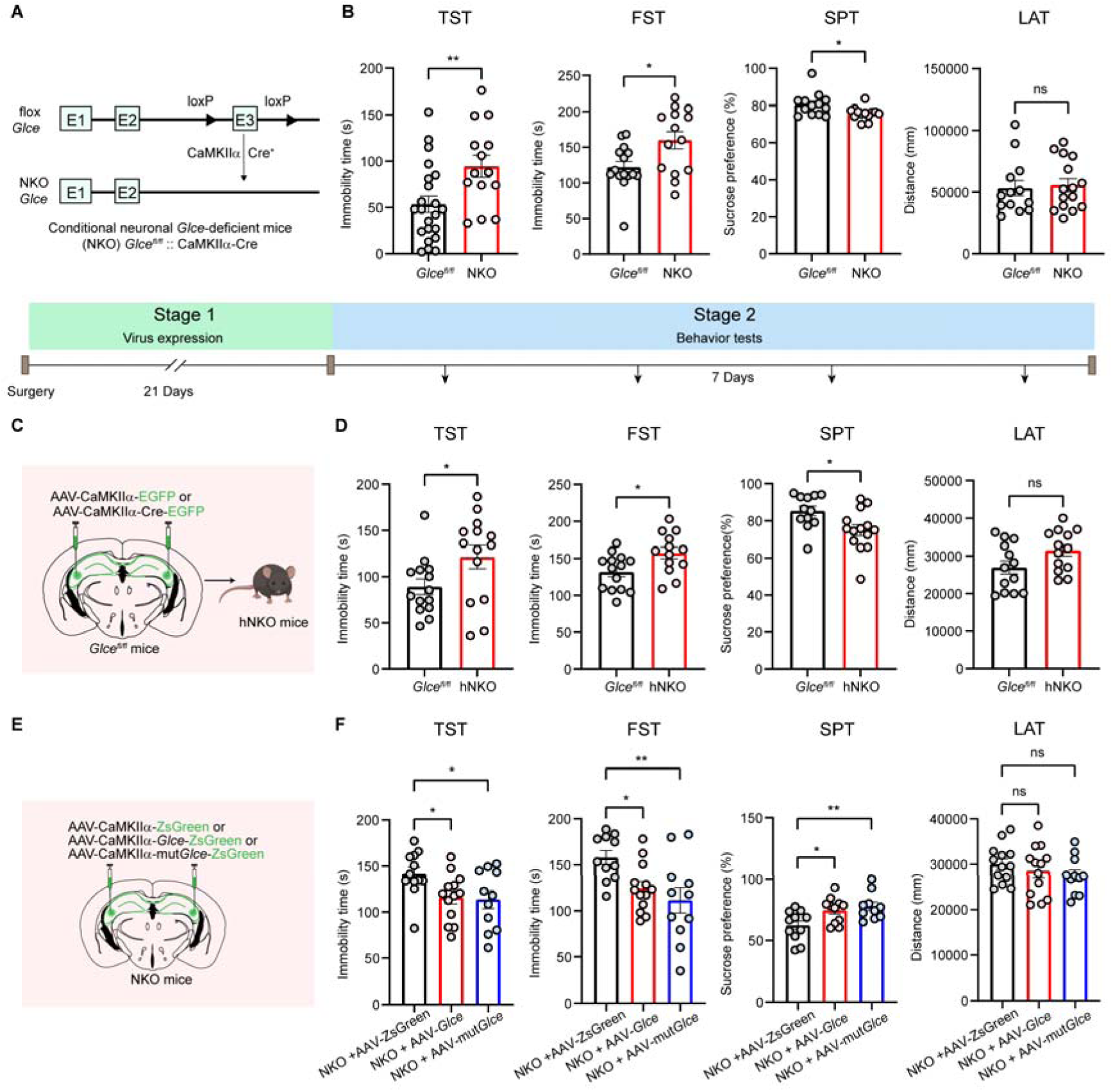
Deletion of *Glce* in the hippocampus leads to depressive-like behaviors, which can be reversed by re-expressing *Glce* in a manner independent of its enzymatic activity. (**A**) Generation of CaMKIIα-Cre mediated *Glce* neuronal-specific knockout (NKO) mice. (**B**) 8- to 12-week-old male NKO mice displayed depressive-like behaviors through TST; FST; SPT and LAT. n = 22, 14 in the TST; n = 15, 14 in the FST; n = 13, 13 in the SPT and n = 13, 15 in the LAT for *Glce^fl/fl^* and NKO group. (**C**) Schematic construction of postnatally neuronal specific-*Glce* knockout in the HPC (hNKO) by AAV in 8- to 12-week-old male *Glce^fl/fl^* mice. (**D**) hNKO mice displayed depressive-like behaviors assessed by TST; FST; SPT and LAT. n = 14, 14 in the TST; n = 14, 13 in the FST; n = 12, 14 in the SPT and n = 13, 13 in the LAT for *Glce^fl/fl^* and hNKO mice. (**E**) Schematic of conditionally *Glce* or *Glce* mutant (mut*Glce*) overexpression in the HPC by AAV of 8- to 12-week-old male NKO mice to identify *Glce* effect in depressive-like behaviors. Here, *Glce* enzymatic mutant sites including Y501F, Y561F and Y579F. (**F**) Effect of *Glce* or *Glce* mutant overexpression on depression-related behaviors in NKO mice represented by TST; FST; SPT and LAT. n = 12, 13, 11 in the TST; n = 11, 13, 11 in the FST; n = 11, 11, 10 in the SPT and n = 13, 13, 11 in the LAT for NKO + AAV-ZsGreen, NKO + AAV-*Glce* and NKO + AAV-mut*Glce* group, respectively. Error bars show s.e.m. Two-tailed unpaired *t* test or Mann-Whitney test (**B** and **D**) or one-way ANOVA, followed by Dunnett’s multiple comparisons test (**F**) was used. ns, not significant, *P < 0.05, **P < 0.01.

To further investigate the specific role of *Glce* in the hippocampus, we firstly used AAV-Syn-*Glce* shRNA to knock down *Glce* expression in the hippocampus (Supplemental Figure 3A). It was found that the expression level of *Glce* was reduced (Supplemental Figure 3B), while no significant depressive-like behaviors were found in mice, characterized by TST, FST, SPT and LAT (Supplemental Figure 3C). We speculated it may be ascribed to heterogeneity across neuronal subpopulations within the hippocampus or incomplete knockdown efficiency of *Glce*. We therefore employed a targeted approach by microinjecting AAV-CaMKIIα-Cre into the hippocampus of *Glce^fl/fl^* mice, creating hNKO mice (Figure 1C and Supplemental Figure 2D). Consistent with our previous observation, these hNKO mice similarly exhibited depressive-like behaviors (Figure 1D).

To further confirm the critical involvement of *Glce* in the hippocampus, we investigated whether re-expression of *Glce* in the hippocampus could reverse the depressive-like phenotypes in NKO mice. As shown in Figure 1, E and F, bilateral infusion of AAV-CaMKIIα-*Glc*e-ZsGreen into the hippocampus of NKO mice significantly alleviated the depressive-like behaviors of NKO mice, as evidenced by reduced immobility in both the TST and FST and restored sucrose preference in the SPT, without altering locomotor. Interestingly, when a non-enzymatic mutant of *Glce* was introduced via AAV-CaMKIIα-mut*Glce*-ZsGreen intra-hippocampal microinjection to inactivate its enzymatic activity (Debarnot et al., 2019; He et al., 2023; Qin et al., 2015), it also alleviated depressive-like behaviors in NKO mice (Figure 1, E and F). These findings suggest that the anti-depressant effects of *Glce* are independent of its enzymatic activity, indicating non-enzymatic role of *Glce* in mood regulation. Taken together, these data support the importance of *Glce* in hippocampal glutamatergic neurons in modulating depressive-like behaviors, independent of its enzymatic function.

### *Glce* facilitates dendritic extension and dendritic outgrowth

In the context of depression, a well-documented observation is the reduced volume and altered structural integrity of the hippocampus, often accompanied by neuronal atrophy, loss and dysfunction (Duman et al., 2016). Given the involvement of *Glce* in stress-induced depressive-like behaviors, we hypothesized that *Glce* might modulate neuronal growth. To assess these effects, we firstly utilized Golgi staining to examine the structure of hippocampal pyramidal neurons in NKO mice. These mice exhibited a significant reduction in dendritic outgrowth (Figure 2A). Notably, an increase in *Glce* levels, whether in its normal or mutant forms, resulted in improved neuronal length in NKO mice (Figure 2B). To further test this, we transfected primary hippocampal neurons with *Glce* knockdown (sh*Glce*) and *Glce* overexpression (*Glce*-OE) lentiviruses simultaneously. We found that co-transfection with a *Glce* overexpression lentivirus in the sh*Glce* group effectively reversed this inhibition, promoting dendritic outgrowth (Supplemental Figure 4). These results indicate the crucial role of *Glce* in maintaining neuronal complexity for resilience against depression.

**Figure 2.**
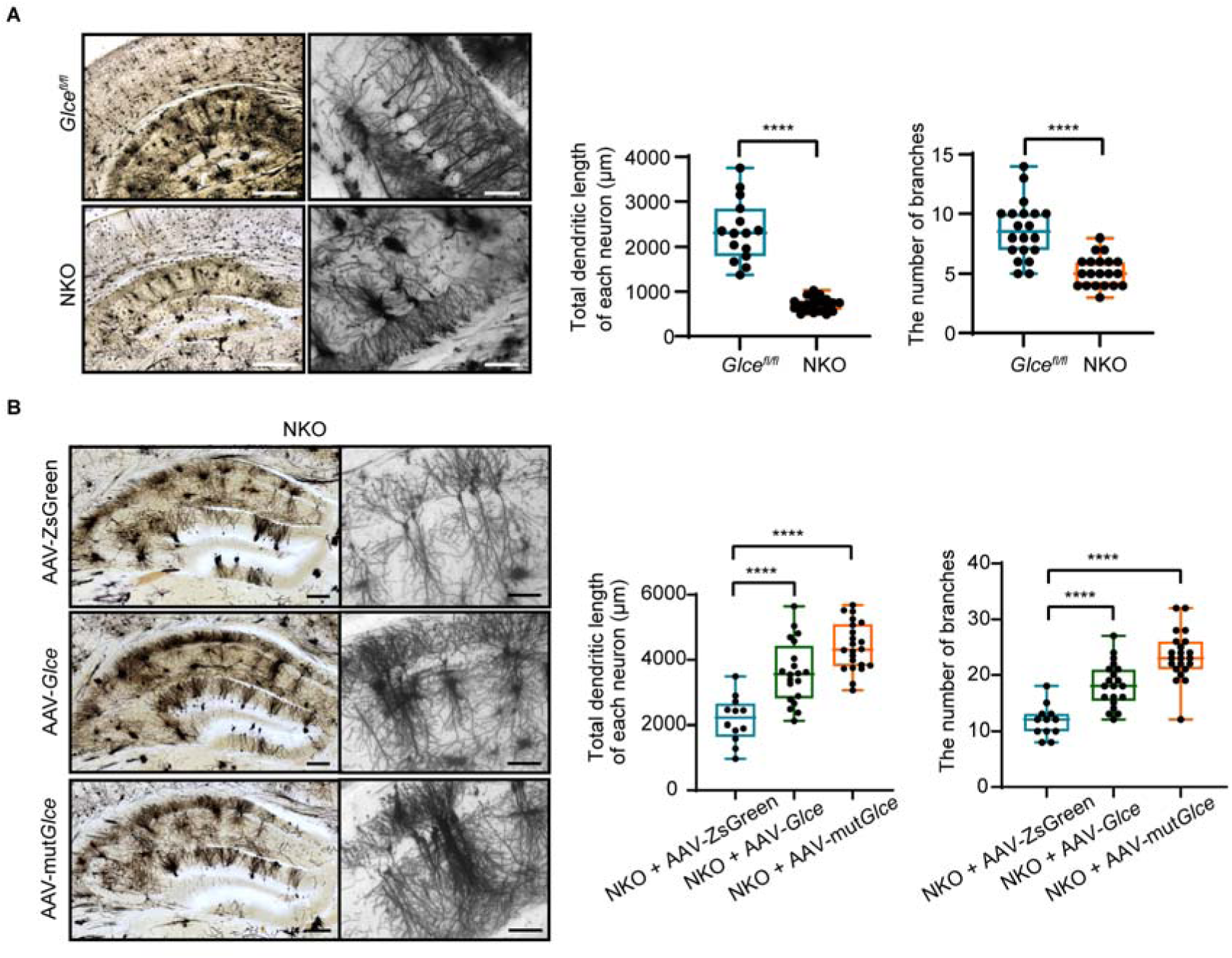
*Glce* promotes dendritic extension and dendritic outgrowth. (**A**) Golgi staining on 100 μm brain slices from NKO and *Glce^fl/fl^* mice. Quantification of dendritic outgrowth employed indicators including total dendritic length and the number of branches of each neuron. Scale bars, 500 μm (left) and 100 μm (right). (**B**) Golgi staining of hippocampal pyramidal neurons on AAV-ZsGreen, AAV-*Glce*, and AAV-mut*Glce* virus injected NKO mice with quantification of dendritic outgrowth. Scale bars, 150 μm (left) and 200 μm (right). Error bars show s.e.m. Two-tailed unpaired *t* test (**A**) or one-way ANOVA, followed by Dunnett’s multiple comparisons test (**B**) was used. ****P < 0.0001.

### *Glce* promotes dendritic outgrowth and ameliorates depressive-like behaviors through the regulation of BDNF expression

BDNF is well recognized for its crucial role in neuroplasticity and promoting dendritic outgrowth in the hippocampus (Duman et al., 2016; Lu et al., 2013; Minichiello, 2009; Wang et al., 2022), as well as its involvement in mediating antidepressant effects and depression treatment (Martinowich et al., 2007; Mizui et al., 2015). We next investigated whether *Glce* might regulate dendritic outgrowth by regulation of BDNF. We first detected BDNF expression in the hippocampus of NKO mice, using immunofluorescence and western blot assays. Interestingly, BNDF level was significantly decreased in the hippocampus of *Glce* deficiency mice, especially in CA3 region (Figure 3, A and B and Supplemental Figure 5). BDNF binds to its high-affinity receptor TrKB, triggering its phosphorylation, and initiates downstream signaling pathways to promoting dendritic outgrowth, synaptic strength and plasticity, with implicated in various neurological and psychiatric disorders (Duman et al., 2016; Minichiello, 2009; Wang et al., 2022). We therefore detected TrKB phosphorylation level, and observed that *Glce* loss significantly impaired TrKB phosphorylation in the hippocampus of NKO mice (Figure 3C). Similarly, hNKO mice also showed reduced levels of BDNF and TrKB phosphorylation in the hippocampus (Figure 3D). Importantly, we found that both *Glce* and mutant *Glce* overexpression increased BDNF expression and enhanced TrKB phosphorylation in NKO mice, supporting that BNDF expression and activity might be modulated by both forms of *Glce* (Figure 3E).

**Figure 3.**
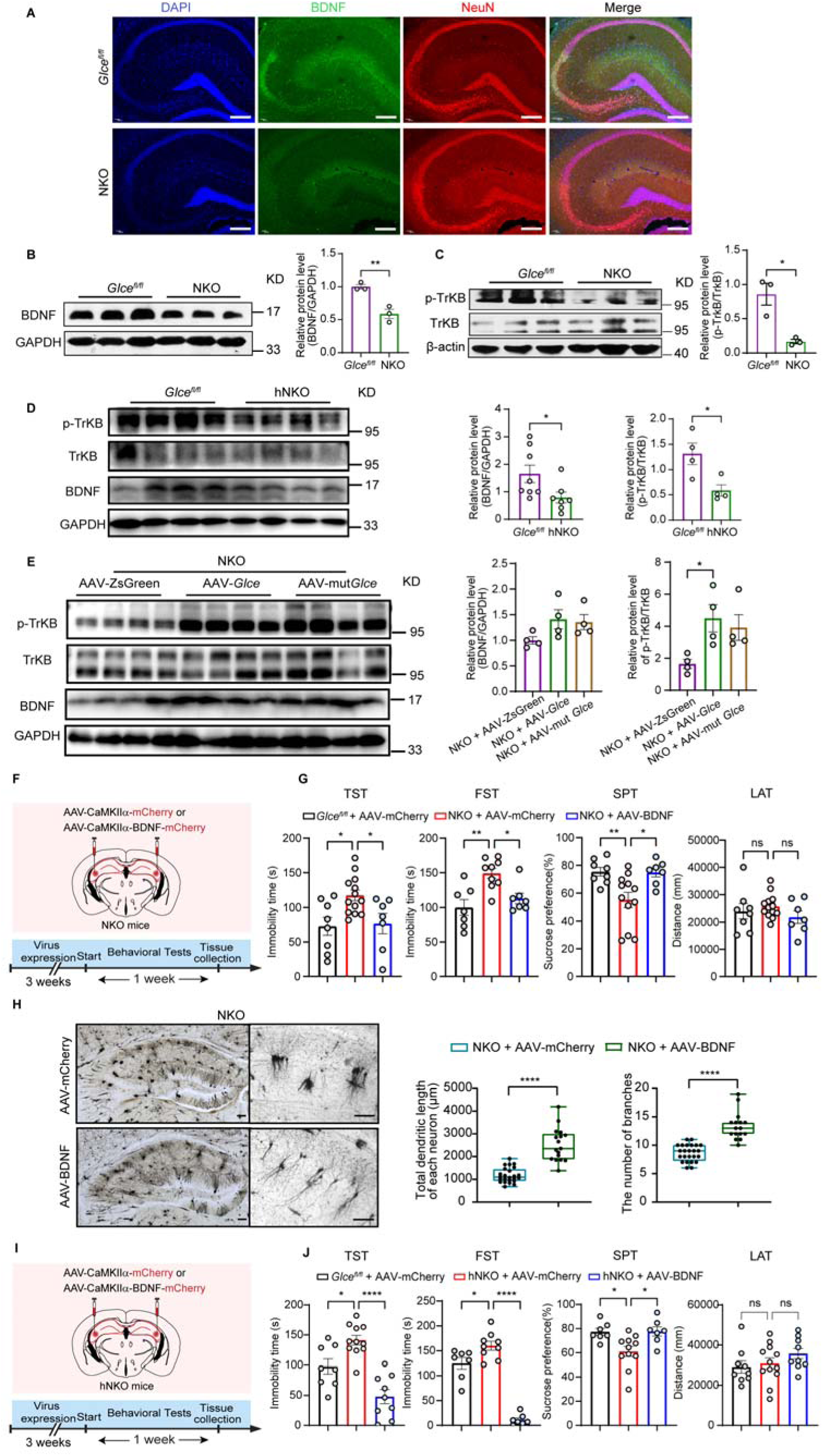
*Glce* facilitates dendritic extension and dendritic outgrowth via enhancing BDNF expression and activity. (**A**) Immunofluorescence images showing BDNF (green) expression in hippocampus of NKO mice and littermate *Glce^fl/fl^* mice. Scale bars, 200 μm. (**B** and **C**) Western blot showing BDNF (B) and p-TrKB/TrKB (C) level in hippocampus of NKO mice, (*Glce^fl/fl^*n = 3, NKO n = 3) with quantification. (**D** and **E**) Effect of *Glce* on BDNF expression and BDNF activity reflected by p-TrKB/TrKB level in hNKO mice (**D**) and adenovirus-mediated *Glce* and mut*Glce* overexpressed NKO mice (**E**) with quantification. (**F** to **J**) Experimental paradigm of BDNF overexpression by AAV, and TST; FST; SPT and LAT were employed to identify BDNF effect on depression-related behaviors in NKO mice (**F**) and hNKO mice (**I**). n = 8, 13, 7 in the TST; n = 7, 9, 7 in the FST; n = 8, 12, 7 in the SPT and n = 8, 13, 7 in the LAT for *Glce^fl/fl^* + AAV-mCherry, NKO + AAV-mCherry and NKO + AAV-BDNF group (**G**). Golgi staining was applied for morphology of hippocampal pyramidal neurons (**H**). Behavioral tests were performed using TST; FST; SPT and LAT to evaluate BDNF function in hNKO mice. n = 8, 11, 9 in the TST; n = 7, 9, 7 in the FST; n = 7, 11, 7 in the SPT and n = 9, 12, 9 in the LAT for *Glce^fl/fl^* + AAV-mCherry, hNKO + AAV-mCherry and hNKO + AAV-BDNF group (**J**). Scale bars, 150 μm and 200 μm (magnified). Error bars show s.e.m. Two-tailed unpaired *t* test (**B** to **D** and **H**) or one-way ANOVA, followed by Dunnett’s multiple comparisons test (**E** and **G** and **J**) was used. ns, not significant, *P < 0.05, **P < 0.01, ****P < 0.0001.

To further determine whether *Glce* deficiency-induced depressive-like behaviors were mediated through BNDF, we bilaterally injected AAV-CaMKIIα-BDNF into the hippocampus of NKO and hNKO mice, followed by behavioral tests (Figure 3, F and I). Indeed, BDNF overexpression significantly reduced immobility in the TST and FST, and increased sucrose preference in the SPT, without altering mice locomotor activity in both NKO and hNKO mice (Figure 3, G and J). Moreover, BDNF markedly improved dendritic outgrowth in these mice (Figure 3H) and retrieved primary hippocampal neuronal growth inhibited by *Glce* knockdown (Supplemental Figure 4). These data indicate that BDNF overexpression can rescue the depressive-like phenotypes in *Glce* deficient mice. Collectively, the data supports that *Glce* facilitates dendritic outgrowth and ameliorates depressive-like behaviors, possibly by regulation of BDNF.

### Modulation of BDNF expression by *Glce* is mediated by potentiation of PI3K/Akt/CREB signaling pathway

BDNF binds and activates TrKB, triggering a cascade of intracellular signaling pathways, among which, the PI3K/AKT signaling pathway is particularly critical for promoting neuronal growth (Duman et al., 2016; Minichiello, 2009; Wang et al., 2022). To investigate the effects of *Glce* on PI3K/AKT signaling, molecular docking analyses were conducted. The results showed that *Glce* might interact with PI3K, particularly forming a complex with the p110α-p85α subunits of hPI3K, however the binding site of which is distinct from the enzymatic active site of *Glce* (Figure 4A). The interaction was further validated by surface plasmon resonance (SPR) studies, which showed that h*Glce* bound to hPI3Kα very strongly with a Kd value of approximately 5.722 × 10^-9^ (Figure 4B). To further confirm the results *in vivo*, co-immunoprecipitation (Co-IP) experiments were carried out. Indeed, *Glce* interacted with PI3Kα in both the primary cortical neurons and mouse hippocampal tissues (Figure 4C). Interestingly, we also found that *Glce* and PI3Kα were colocalized in the Golgi apparatus (Figure 4D), aligning with previous findings (Crawford et al., 2001).

**Figure 4.**
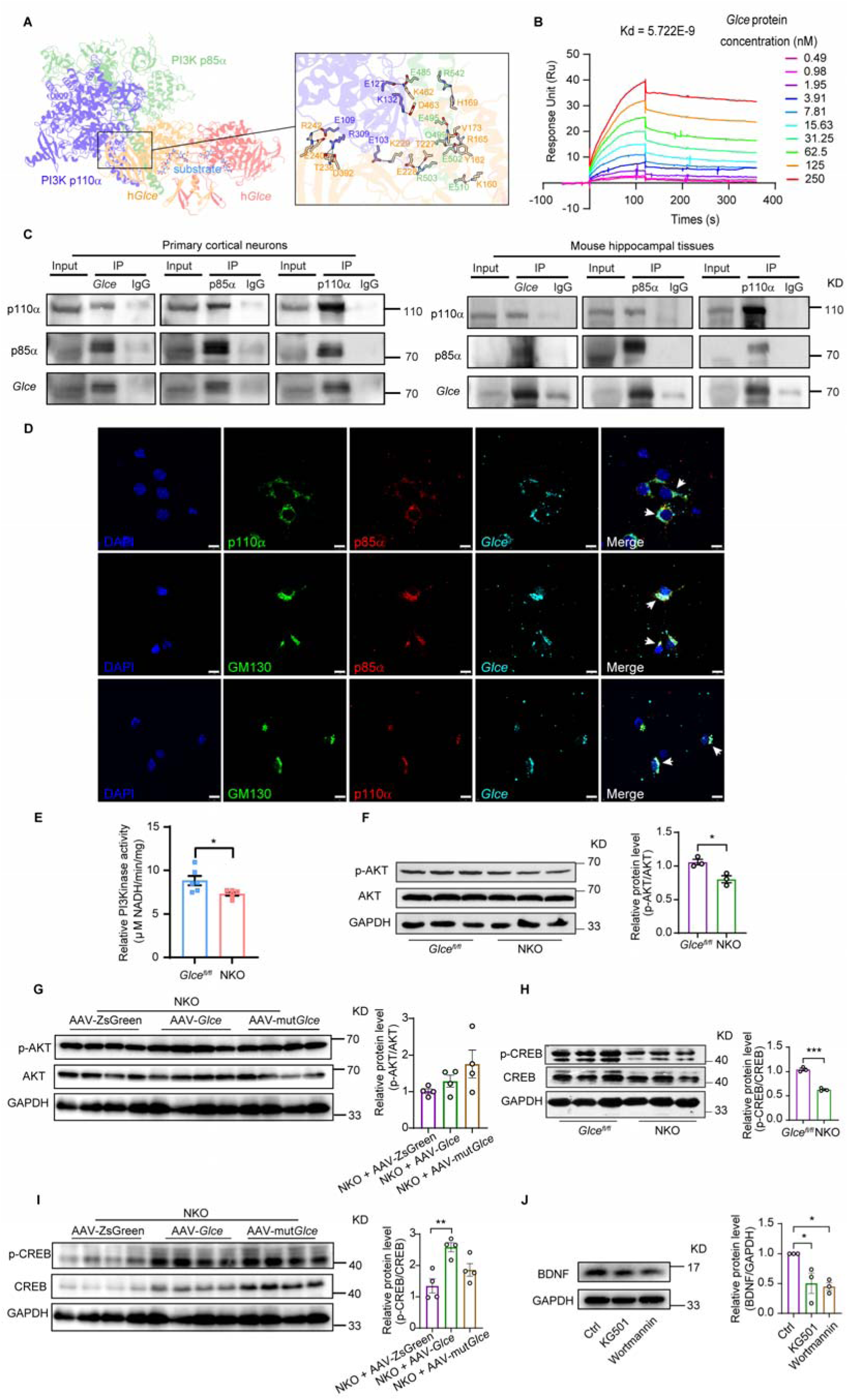
*Glce* enhances BDNF expression and activity via facilitating PI3K/Akt/CREB signaling pathway. (**A**) The docking model of the hPI3K-h*Glce* complex. (**B**) Direct binding between *Glce* and PI3Kα (p85α and p110α) proteins was measured by SPR. (**C**) Co-immunoprecipitation (Co-IP) showing the interaction between *Glce* and PI3Kα protein in primary cortical neurons and in hippocampal tissues by using anti-*Glce*, anti-PI3K p85α and p110α antibodies bidirectional pull-down. (**D**) PI3Kα co-localized with *Glce* in the Golgi apparatus in primary hippocampal neurons. Scale bars, 25 μm. (**E**) Relative PI3Kinase activity was examined in hippocampus tissues of NKO mice, n = 6 mice per group. (**F** and **G**) Western blot analysis of p-AKT/AKT protein level in the hippocampus tissues of NKO mice (**F**) and *Glce* or mut*Glce* overexpressed NKO mice (**G**) versus respective *Glce^fl/fl^*mice with quantification. (**H** and **I**) Western blot (left) and quantification (right) showing the effect of *Glce* on CREB activation represented by p-CREB/CREB in the hippocampus of NKO mice (**H**) and *Glce* or mut*Glce* overexpressed NKO mice (**I**). (**J**) BDNF protein level was detected after treatment of KG501 (10 μM) and wortmannin (1 μM) in the primary cortical neurons (DIV3) from C57BL/6J mice for 24 hours by Western blot (left) with quantification (right). Error bars show s.e.m. Two-tailed unpaired *t* test (**E** and **F** and **H**) or one-way ANOVA, followed by Dunnett’s multiple comparisons test (**G** and **I** and **J**) was used. *P < 0.05, **P < 0.01, ***P < 0.001. Note: Figure 3E, Figure 4G and I were derived from the same Western blot experiment with common internal control GAPDH band.

To investigate the role of *Glce* in modulating PI3K, we detected whether *Glce* regulated the PI3K enzymatic (PI3Kinase) activity and downstream AKT phosphorylation. PI3K activity assay showed a reduced PI3K activity within the hippocampus of NKO mice (Figure 4E). This decrease in PI3K activity was accompanied by reduced level of AKT phosphorylation in the hippocampus of NKO mice (Figure 4F). The PI3K/AKT pathway is known to regulate CREB phosphorylation (Marsden, 2013; Nestler et al., 2002), a process that stimulates gene transcription such as BDNF, thereby promoting dendritic outgrowth and establishing a positive feedback loop (Chen et al., 2012; Koo et al., 2015; Wang et al., 2022). In line with this, the CREB phosphorylation level was reduced in the hippocampus of NKO mice (Figure 4H). Similarly, hNKO mice also displayed decreased phosphorylation levels of AKT and CREB in the hippocampus (Supplemental Figure 6). Moreover, overexpression of *Glce*, as well as mutant *Glce*, might significantly activate AKT and CREB phosphorylation in NKO mice (Figure 4, G and I), supporting the role of *Glce* in modulating this signaling pathway. Finally, to confirm the involvement of PI3K-AKT-CREB signaling pathway, KG501, a selective CREB inhibitor, and wortmannin, a selective PI3K inhibitor were employed. We found that disruption of the PI3K-CREB signaling pathway in primary cortical neurons resulted in a significant reduction in BDNF expression (Figure 4J). Collectively, these findings suggest that *Glce* regulates BDNF via the PI3K-AKT-CREB signaling pathway.

### *Glce* regulates depressive-like behaviors via targeting and activating PI3K

Next, we asked whether PI3K served as a key downstream molecule underlying *Glce* deficit-mediated dendritic growth impairment and depressive-like behaviors. To explore this possibility, PI3K inhibitor LY294002 was bilaterally infused into the mice hippocampus, followed by depressive-like behavior tests (Figure 5A). The results showed that inhibition of PI3K activity in the hippocampus significantly induced depressive-like behaviors (Figure 5B) and suppressed dendritic outgrowth (Figure 5C), suggesting that PI3K inhibition contributes to depressive-like behaviors, possibly through reduced dendritic extension.

**Figure 5.**
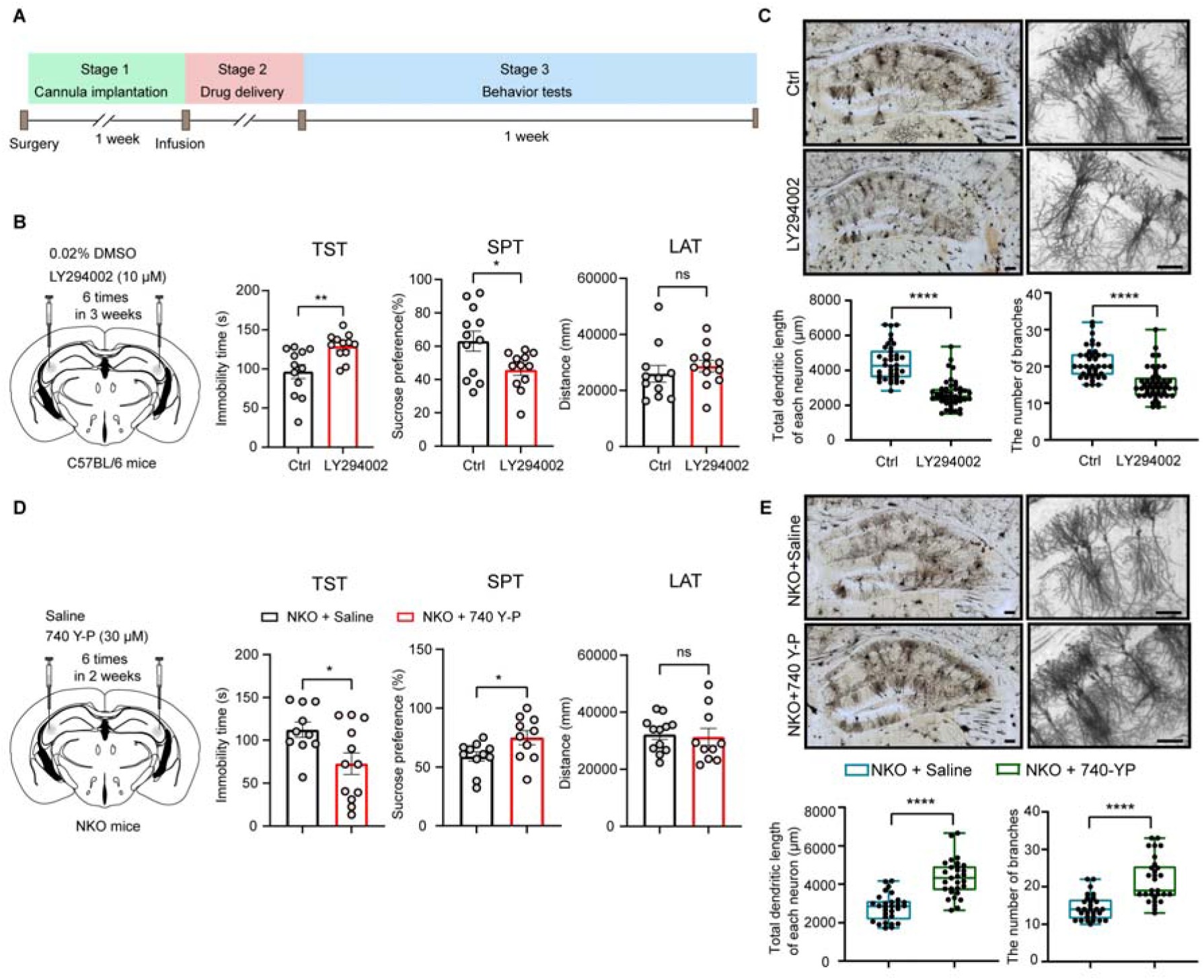
*Glce* exerts its effects on depressive-like behaviors mediated by PI3K. (**A**) Schematic of stereotactic brain drug administration paradigm. (**B** and **C**) Bilateral infusion of LY294002 (10 μM) and 0.02% DMSO as solvent control via cannulae implantation in the HPC of 8-week-old male C57BL/6J mice, with dosing interval of 6 times in 3 weeks and then TST, SPT and LAT were employed to identify the effect of LY294002 on depressive-like behaviors. n = 12, 12 in the TST; n = 12, 12 in the SPT and n = 11, 12 in the LAT for Ctrl and LY294002 group, respectively. LY294002, a PI3K specific selective inhibitor (**B**). Golgi staining on brain slices from mice with LY294002 and 0.02% DMSO, with qualification of dendritic outgrowth (5-6 neurons per mouse). (**C**). Scale bars, 200 μm. (**D** and **E**) Bilateral infusion of 740 Y-P (30 μM) and saline as solvent control via cannulae implantation in the HPC of 8- to 12-week-old male NKO mice with dosing interval of 6 times in 2 weeks. Then TST; SPT and LAT were employed to identify the effect of 740 Y-P on NKO mice. n = 10, 12 in the TST; n = 11, 10 in the SPT and n = 12, 10 in the LAT for saline and 740 Y-P group, respectively. 740 Y-P, a PI3K specific selective agonist (**D**). Golgi staining on brain slices from NKO mice administrated with saline and 740 Y-P, with qualification of dendritic outgrowth. (**E**). Scale bars, 200 μm. Error bars show s.e.m. Two-tailed unpaired *t* test or Mann-Whitney test (**B** to **E**) were used. ns, not significant, *P < 0.05, **P < 0.01, ****P < 0.0001.

We then explored whether activation of PI3K could rescue the depressive-like phenotypes induced by *Glce* deficiency. To examine, PI3K agonist 740 Y-P was bilaterally injected into the hippocampus of NKO mice and followed by behavioral tests. Indeed, 740Y-P ameliorated depressive-like behaviors and dendritic growth suppression caused by *Glce* deficiency (Figure 5, D and E). These data suggest that PI3K may serve as a key downstream molecule underlying *Glce*-regulated dendritic growth and depressive-like behaviors.

### Chronic stress impairs hippocampal *Glce* expression, impedes dendritic outgrowth and induces depressive-like behaviors

Chronic stress is a prominent contributor to the development of major depressive disorder (MDD). Since *Glce* is involved in depressive-like behaviors, we sought to determine whether *Glce* expression was affected by chronic stress. We first generated two models: a ten-day chronic social defeat stress model (CSDS) and a three-week chronic restrained stress model (CRS) (Figure 6A and Supplemental Figure 7A). As shown in Figure 6, B to D, CSDS mice demonstrated significant social avoidance, increased immobility in the FST and decreased sucrose preference in the SPT. CRS mice also displayed increased immobility in the FST and TST, decreased sucrose preference in the SPT (Supplemental Figure 7, B to D). Neither stress altered mice locomotor activity in the LAT (Figure 6E and Supplemental Figure 7E). Using these models, we detected the expression level of *Glce* in the hippocampus. Surprisingly, chronic stress significantly decreased *Glce* expression (Figure 6F and Supplemental Figure 7F). To validate these observations, serum samples from individuals with depression and healthy controls were collected, followed by the *Glce* expression determination. The results showed a statistically significant decrease in *Glce* concentration in the serum from depression patients (Supplemental Figure 8).

**Figure 6.**
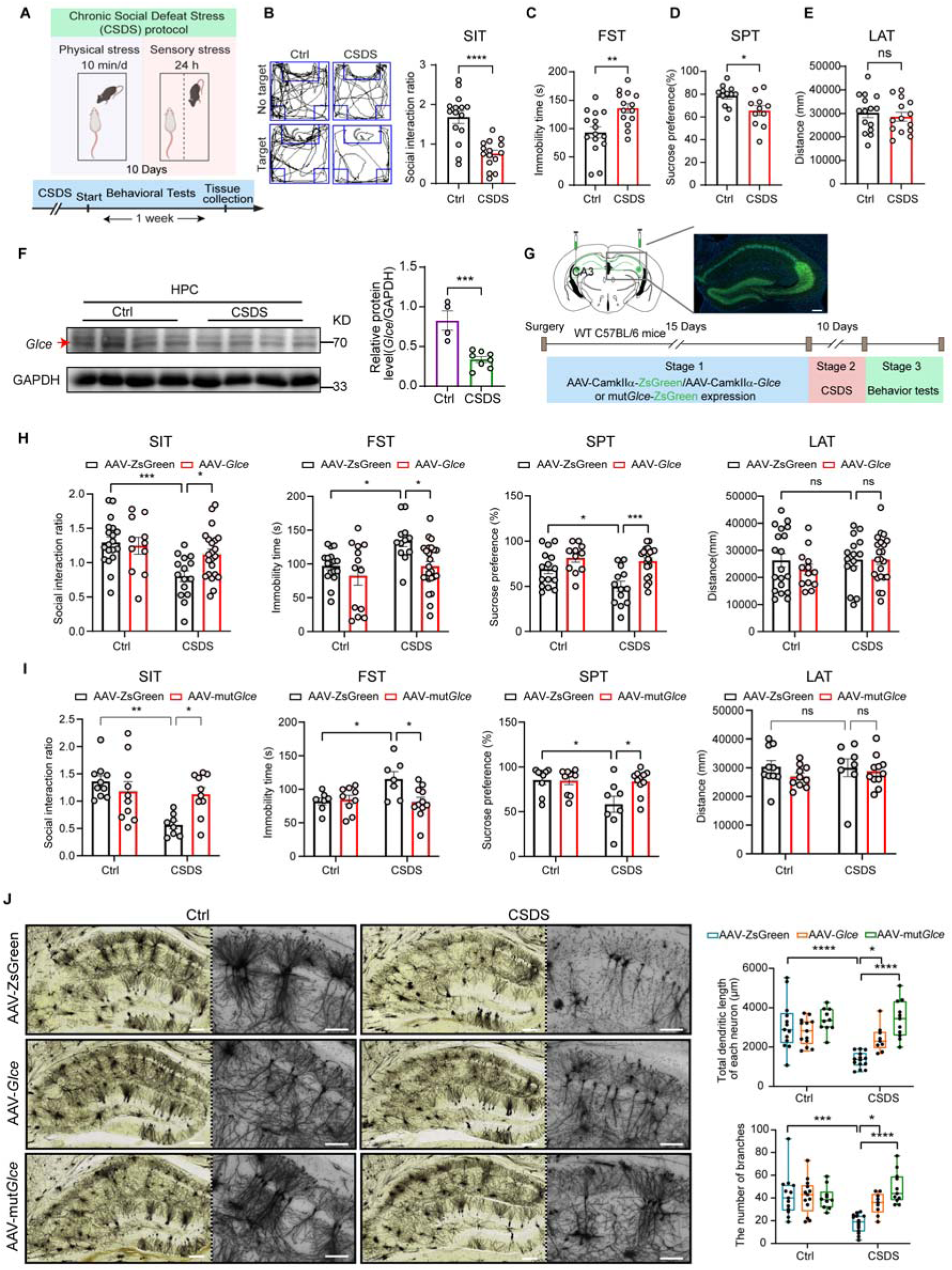
Chronic stress reduces hippocampal *Glce* expression and triggers depressive-like behaviors. (**A**) Schematic of the CSDS paradigm. (**B** to **E**) Behavior tests of CSDS model mice and relative wildtype C57BL/6 Ctrl mice examined by SIT (**B**); FST (**C**); SPT (**D**) and LAT (**E**). n = 15, 15 mice in the SIT; n = 15, 14 mice in the FST; n = 12, 11 mice in the SPT and n = 15, 14 mice in the LAT for Ctrl and CSDS group. (**F**) Western blot showing *Glce* protein expression marked by the red arrow in hippocampus from CSDS model mice with quantification, (Ctrl n = 4, CSDS n = 8). (**G**) Experimental diagram of *Glce* or *Glce* mutant (mut*Glce*) overexpression in the HPC of 8-week-old male C57BL/6J mice by AAV following CSDS. Representative image showing ZsGreen expression in the HPC (upper). (**H** and **I**) Effect of *Glce* (**H**) and mut*Glce* (**I**) overexpression with or without CSDS stimulus as assessed by SIT; FST; SPT and LAT. N = 19, 11, 15, 22 in the SIT; n = 17, 13, 12, 24 in the FST; n = 15, 11, 13, 20 in the SPT and n = 19, 13, 15, 22 in the LAT for Ctrl + AAV-ZsGreen, Ctrl + AAV-*Glce*, CSDS + AAV-ZsGreen and CSDS + AAV-*Glce* group; n = 10, 10, 8, 10 in the SIT; n = 7, 9, 7, 11 in the FST; n = 9, 10, 8, 11 in the SPT and n = 10, 10, 8, 11 in the LAT for Ctrl + AAV-ZsGreen, Ctrl + AAV-mut*Glce*, CSDS + AAV-ZsGreen and CSDS + AAV-mut*Glce* group, respectively. (**J**) Golgi staining depicting pyramidal neurons in hippocampus of CSDS-treated or untreated *Glce*, *Glce*-mutant and relative Ctrl mice, represented by total dendritic length and the number of branches of each neuron. Scale bar, 200 μm. Error bars show s.e.m. Two-tailed unpaired *t* test (**B** to **F**) or ordinary two-way ANOVA (**H** to **J**), followed by Tukey’s multiple comparisons test was used. ns, not significant, *P < 0.05, **P < 0.01, ***P < 0.001, ****P < 0.0001.

To confirm the critical role of *Glce* in depression, AAV-CaMKIIα-*Glce* was microinjected in the hippocampus to locally overexpress *Glce* (Figure 6G). Indeed, *Glce* overexpression significantly ameliorated CSDS-induced depressive-like behaviors, as evidenced by increased social interaction in the SIT, decreased immobility time in the FST, and increased sucrose preference in the SPT (Figure 6H). Similarly, we found that overexpression of mut*Glce* also reversed CSDS-induced depressive-like behaviors (Figure 6I), also supporting its non-enzymatic function in depression management. Moreover, both *Glce* and mut*Glce* overexpression ameliorated CSDS-induced attenuation of dendritic outgrowth and BDNF expression (Figure 6J and Supplemental Figure 9). Taken together, these data suggest that chronic stress decreases *Glce* expression in the mice hippocampus, contributing to impaired dendritic outgrowth and depressive-like behaviors.

### The downregulation of *Glce* expression is attributed to chronic stress-induced elevation of miR-34c-5p expression

Consistent with the aforementioned findings, we observed significant reductions in *Glce* mRNA levels, PI3K activity and BDNF mRNA levels in the hippocampus of mice subjected to the CSDS (Figure 7, A to C). We next sought to understand the mechanism underlying the chronic stress-induced decrease in *Glce* and subsequent changes.

**Figure 7.**
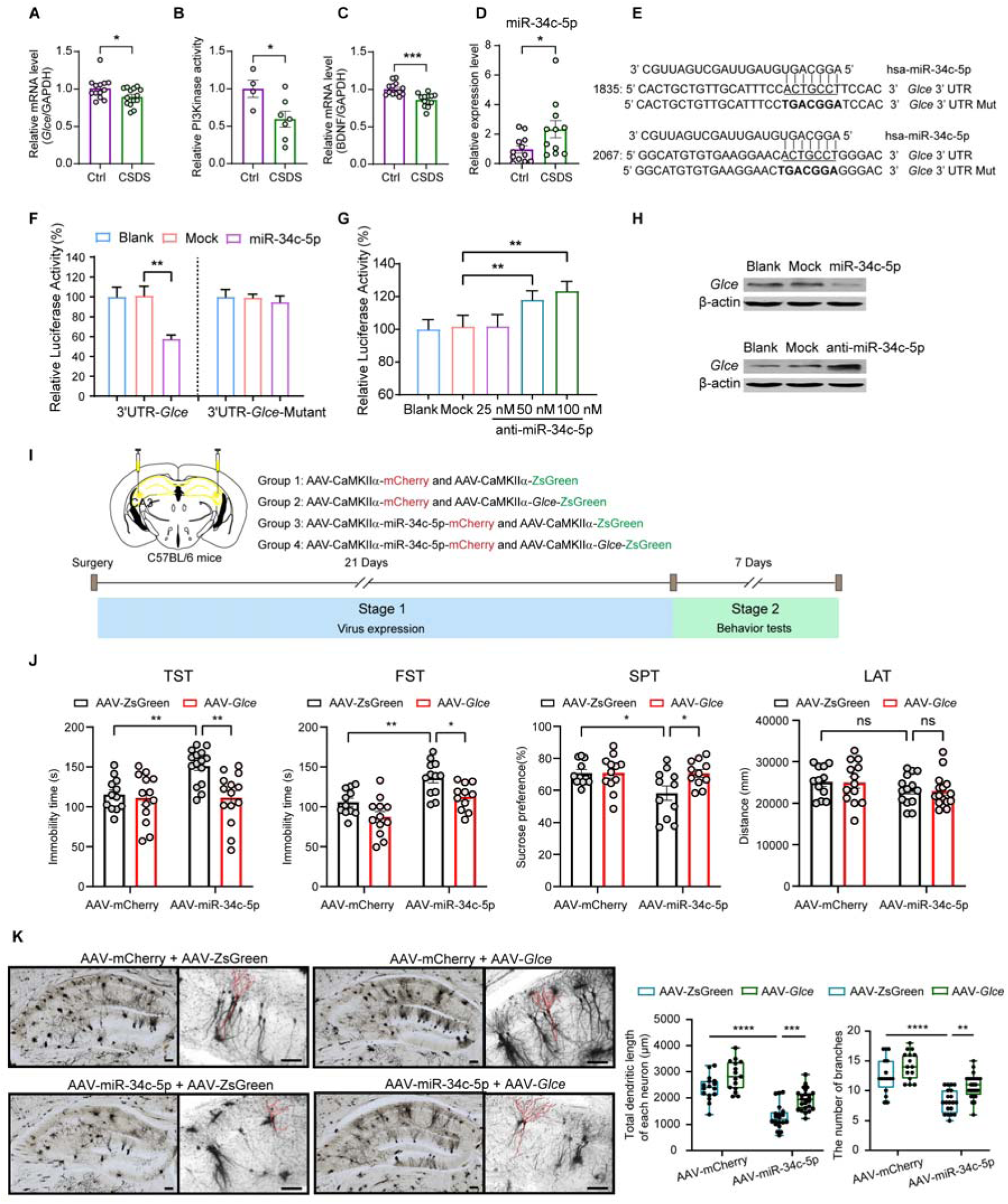
The reduction of *Glce* expression is driven by the upregulation of miR-34c-5p caused by chronic stress. (**A** to **C**) *Glce* mRNA level (Ctrl n = 13, CSDS n = 14) (**A**), PI3Kinase activity (Ctrl n = 4, CSDS n = 7) (**B**) and BDNF mRNA level (Ctrl n = 13, CSDS n = 13) (**C**) were tested by qPCR and PI3Kinase activity detection kit in the hippocampus tissues from mice with or without CSDS stimulus. (**D**) qPCR showing miR-34c-5p levels (Ctrl n = 12, CSDS n = 11) in hippocampus from mice with or without CSDS stimulus. (**E**) Alignment of potential miR-34c-5p binding sites in the 3’UTR of *Glce* mRNA. (**F**) HEK293T cells were co-transfected with miR-34c-5p (miR-34c-5p) or vector control (Mock) plus *Glce* 3’UTR (3’UTR-*Glce*) or mutant-*Glce* 3’UTR (3’UTR-*Glce*-Mutant). Luciferase activities were measured after transfection for 36 h. (**G**) HEK293 cells were co-transfected with *Glce* 3’UTR (3’UTR-*Glce*) and anti-miR-34c-5p at concentrations of 0, 25, 50, or 100 nM for 36 h, followed by performing the luciferase assay. (**H**) Protein level of *Glce* in primary cortical neurons from Sprague-Dawley rat transfected with miR-34c-5p (miR-34c-5p) or vector control (Mock), anti-miR-34c-5p (anti-miR-34c-5p) or the vector control (Mock), repeating at least three times. (**I** to **K**) miR-34c-5p overexpression induced depressive-like phenotypes in C57BL/6J mice which was reversed by *Glce* overexpression. Schematic of experiment procedure including virus brain stereotactic injection and behavioral tests (**I**). Behavioral tests were examined using TST; FST; SPT and LAT. n = 12, 13, 15, 14 in the TST; n = 11, 12, 12, 11 in the FST; n = 11, 12, 11, 11 in the SPT and n = 12, 13, 14, 15 in the LAT for AAV-ZsGreen + AAV-mCherry, AAV-*Glce* + AAV-mCherry, AAV-ZsGreen + AAV-miR-34c-5p and AAV-*Glce* + AAV-miR-34c-5p group (**J**). Golgi staining was applied for morphology of hippocampal pyramidal neurons, represented by the number of branches and total dendritic length of each neuron (4-5 neurons per mouse) (**K**). Scale bars, 150 μm and 200 μm. Error bars show s.e.m. Two-tailed unpaired *t* test or Mann-Whitney test (**A** to **D**) or one-way ANOVA (**F** and **G**) or ordinary two-way ANOVA, followed by Tukey’s multiple comparisons test (**J** and **K**) was used. ns, not significant, *P < 0.05, **P < 0.01, ***P < 0.001, ****P < 0.0001.

MicroRNAs are widely reported to downregulate protein expression and are implicated in major depressive disorders (Fan et al., 2022; Feng et al., 2023; Li et al., 2021). We hypothesized that *Glce* expression might also be regulated by miRNA in CSDS induced mice. Interestingly, we found a significant increase in miR-34c-5p expression in CSDS-induced depression model (Figure 7D), which aligns with previous literature (Yu et al., 2024). This prompted us to explore the specific role of miR-34c-5p in major depressive disorders.

We firstly conducted a bioinformatic analysis, surprisingly finding that miR-34c-5p might bind to the 3’UTR of the *Glce* mRNA, as predicted by the PicTar algorithm (Figure 7E). To validate this prediction, we constructed luciferase reporter vectors encoding the *Glce* 3’UTR and co-transfected them with miR-34c-5p mimics into HEK293T cell. The results showed a significant reduction in luciferase activity (Figure 7F), indicating a direct binding between miR-34-5p and the *Glce* 3’UTR. We further employed site-directed mutagenesis to introduce specific base pair mutations in the *Glce* 3’UTR. When this mutated reporter (*Glce* 3’UTR Mut) was co-transfected with miR-34c-5p, no change in luciferase activity was observed (Figure 7F), confirming that miR-34-5p specifically targets *Glce* 3’UTR. We then generated a 2-O-methyl-modified oligo RNA (anti-miR-34c-5p), designed to complement and anneal to miR-34c-5p. We found that the anti-miR-34-5p significantly block the effects of miR-34-5p, resulting in increased luciferase activity (Figure 7G). Moreover, we observed that miR-34c-5p markedly reduced *Glce* expression, whereas anti-miR-34c-5p enhanced it (Figure 7H). These data support that miR-34c-5p targets and downregulates *Glce*.

We further generated AAV-CaMKIIα-miR-34c-5p-mCherry to selectively upregulate miR-34c-5p level in the hippocampus to assess the role of miR-34c-5p on depression related behaviors. Simultaneously, we introduced *Glce* through AAV-CaMKIIα-*Glce*-ZsGreen to determine if *Glce* supplementation could counteract the behavioral changes elicited by miR-34c-5p overexpression (Figure 7I). The results suggested that overexpression of miR-34c-5p increased immobility time in the TST and FST and decreased sucrose preference in the SPT, without altering mice locomotor activity. These depressive-like behaviors were reversed by *Glce* overexpression (Figure 7J). Golgi staining further revealed that miR-34c-5p significantly impaired dendritic outgrowth, whereas *Glce* supplementation restored it (Figure 7K). Collectively, these findings suggest that miR-34c-5p acts as a key upstream regulator of *Glce*, modulating both dendritic outgrowth and depressive-like behaviors.

## Discussion

Dendrite arbor morphology is pivotal for neural connectivity, synaptic structure plasticity and brain function (Copf, 2016; Emoto, 2011). The initial dendritic arbor morphology is established early during embryonic development; however, arbors are dynamic and retain their gross morphology by employing active maintenance mechanisms throughout the adult life. Irregularities in either the early developmental phase, or in the later maintenance phase will lead to either neurodevelopmental disorders such as autism, Down’ and Rett syndromes (Armstrong et al., 1995; Becker et al., 1986; Kaufmann and Moser, 2000; Pardo and Eberhart, 2007; Walsh et al., 2008), or neuropsychiatric diseases such as schizophrenia, anxiety and depression (Bellon et al., 2011; Cotter et al., 2000; Eiland and McEwen, 2012; Kulkarni and Firestein, 2012; Soetanto et al., 2010). Nevertheless, the molecular mechanisms underlying the maintenance of dendritic morphology are poorly understood. In this study, we identify that *Glce* is critical for maintaining normal dendritic morphology. Knockout (KO) of *Glce* in hippocampal neurons prevents dendrite outgrowth and induces dendrite atrophy. Reintroduction of *Glce* into the hippocampus of *Glce* KO mice rescues dendrite atrophy. Although substantial evidence has demonstrated the critical role of HSPGs and their correct modification by HS-modifying enzymes in neuron development and neuropsychiatric diseases (Irie et al., 2012; Li et al., 2002; Zhang et al., 2018), we surprisingly find that the facilitatory effect of *Glce* on dendrite outgrowth and the resistant effect of *Glce* on depression is independent on its enzymatic activity of HS-modification, since mutations of *Glce* enzymatic active sites fails to changes in dendrite outgrowth.

One of the most important traits of neurons is the capacity to dynamically adapt to changes in neuronal activity and environmental stimuli. Dendrites, comprised of dendritic arbors and spines, are cellular elements that are responsible for this kind of plasticity. While the causative relationship between dendritic spine abnormalities and neurodevelopmental and neuropsychiatric disorders is well established, such relationship remains elusive for dendritic arborization impairments. Dendrite arbors also display significant remodeling in response to neuronal activity and extrinsic stimuli (Miller and Kaplan, 2003; Wong and Ghosh, 2002). For instance, chronic stress, a major predisposing factor for depression, reduces dendritic length, branching and complexity and results in dendrite atrophy in the rodent hippocampus and prefrontal cortex (PFC) (Li et al., 2011; Liu and Aghajanian, 2008; Magariños et al., 1996; McEwen et al., 2012; Morrison and Baxter, 2012; Radley et al., 2004). Consistently, reduced volume of the hippocampus and PFC is also observed in people with major depressive disorders (MDD) (Huang et al., 2013; Price and Drevets, 2010; Ressler and Mayberg, 2007; Shin and Liberzon, 2010). These studies suggest that dendrite atrophies are linked to develop depressive disorders and point out that the maintenance of normality of dendritic outgrowth is important for brain function. However, the molecular mechanisms and signaling pathway by which chronic stress results in abnormal dendrite outgrowth are poorly understood. This study provides evidence to show *Glce* acting as a crucial regulator for dendrite outgrowth and reveals that downregulation of *Glce* levels, as a result of upregulation of miR-34c-5p expression, contributes to chronic stress-induced changes in dendritic outgrowth.

Phosphatidylinositol 3-kinase (PI3K) is important for both dendritic arborization and morphogenesis (Cuesto et al., 2011; Hood et al., 2020; Jaworski et al., 2005; Kumar et al., 2005; Sánchez-Castillo et al., 2022). Akt is an important effector through which PI3K controls dendritic morphogenesis (Jaworski et al., 2005; Kumar et al., 2005; Lim and Walikonis, 2008; Urbanska et al., 2012). Moreover, Akt has been demonstrated to regulate growth factors BDNF and HGF-induced dendrite morphogenesis in cultured hippocampal neurons (Lim and Walikonis, 2008; Zheng et al., 2008). In present study, we demonstrate that the regulatory effect of *Glce* on dendritic morphology is dependent on its molecular scaffold function that directly engages the PI3K to drive a downstream Akt/CREB/BDNF pathway, rather than its HS-modifying activity. we uncover that PI3K can binding to *Glce* via its regulatory subunit p85α and catalytic subunit p110α, through which *Glce* controls dendrite outgrowth. This is manifested by the findings that intra-hippocampus administration of PI3K inhibitor significantly reduces dendrite outgrowth. In contrast, intra-hippocampus administration of PI3K agonist rescues dendrite outgrowth of KO mice. Moreover, pharmacological manipulations of PI3K activity with PI3K inhibitor and agonist reversely regulate depressive-like behaviors.

BDNF plays an essential role for synaptic development and plasticity. BDNF can regulate synapse formation through increasing the arborization of dendrites (Luikart et al., 2005; Park and Poo, 2013; Vicario-Abejón et al., 2002). Mice with BDNF deletion or with a knock-in of a human loss-of-function BDNF gene variant (Val66Met) displays reduction of dendrite branching and spine density in the hippocampus (Chen et al., 2006; Chiaruttini et al., 2009; Liu et al., 2012; Magariños et al., 2011; Yu et al., 2012). Moreover, there is also evidence that BDNF levels are decreased in postmortem brains of depressed patients (Duman and Voleti, 2012; Krishnan and Nestler, 2008) and targeted deletion of BDNF in the hippocampus is sufficient to cause depressive behaviors by stress (Duman and Voleti, 2012; Yu et al., 2012). In consistent with these studies, the results of our study support a key role of BDNF in modulation of dendrite morphology and depressive behaviors and demonstrates that *Glce* can control BDNF levels in the hippocampus through PI3K/Akt/CREB pathway. Our data suggests that in the process of changes in BDNF expression and dendritic morphology initiated by chronic stress, BDNF and PI3K act as mutual effectors. Chronic stress decreases hippocampal BDNF expression and induces dendritic atrophies through a negative feedback cycle initiated by *Glce*, in which chronic stress downregulates *Glce* expression through upregulation of miR-34c-5p and resultantly impairs PI3K activity, leading to decreased BDNF expression as result of attenuation of PI3K/Akt/CREB pathway signaling. Reduction of BDNF levels would further impair PI3K activity as substantial evidence points out that PI3K is a critical effector of BDNF (Wang et al., 2022).

This study still has several limitations. One is that we did not establish whether *Glce* levels were also decreased in the postmortem hippocampus of human subjects with depression. Although we found that decreased *Glce* levels were occurred in the plasma of human subjects with depression relative to matched controls, it may not represent same changes occurred in hippocampus of human subjects with depression. Future research should confirm that this change was also occurred in hippocampus of human subjects with depression. If it is the case, *Glce* may be a potential biomarker for depression diagnosis. The second limitation is that lack of use of females in our experiments. Future studies should include female animals to comprehensively investigate the role of *Glce* defect in chronic stress-induced depression and to uncover potential sex differences, which will help enhance the generalizability and clinical relevance of the research findings. Additionally, as an enzyme integral to the modification of heparan sulfate, *Glce* is pivotal in maintaining the structural integrity and endowing the diverse biological functions of heparan sulfate proteoglycans. Therefore, elucidating the molecular underpinnings of how alterations in HSPG biology, consequent to *Glce* deficiency, contribute to the pathogenesis of depressive symptoms is a compelling avenue for future research. Moreover, while behavioral tests such as the TST and FST are widely used to assess despair-like behaviors in mouse models of depression-like behaviors, they present limitations related to animal welfare concerns and challenges in result interpretation. Thus, the methodological approach to detecting depression-related behaviors in mice itself constitutes a limitation.

In sum, this study identifies *Glce* as a key regulator for dendritic complexity and establishes that its regulatory is mediated through a non-canonical signaling role, rather than its HS-modifying activity. Glce acts as a molecular scaffold that directly binds to and activates the PI3K complex, subsequently upregulating BDNF expression to maintain dendritic integrity.

## Methods

### Animals

All animal procedures were conducted and approved in compliance with the guidelines and protocols (2022-06-DK-107) by the Institution Animal Care and Use Committee at Shanghai Institute of Materia Medica. Mice were maintained in the core animal facility and housed under a 12 h light/dark cycle with ad libitum access to food and water. Food and water were temporarily restricted only when necessary for specific experimental procedures. The mice used in the experiments were male C57BL/6J, aged 8 or 12 weeks. For primary neurons cultures, the female C57BL/6J mice or Sprague-Dawley rat at the embryonic day 16 were sourced from Shanghai SLAC Laboratory Animal (Shanghai, China).

The floxed *Glce* mice and CaMKIIa-Cre transgenic mice were obtained from Shanghai Model Organisms (Shanghai, China). In the floxed *Glce* mice, loxP sequences flanked exon 3 of the *Glce* gene (Li et al., 2003). To create neuronal-specific *Glce* knockout mice, *Glce^flox/flox^*mice were crossed with CaMKIIα-Cre transgenic mice to delete *Glce* exon 3. Genotyping of *Glce* knockout mice were performed using PCR with designed primers including: loxP site of the targeted *Glce* allele: forward, 5’-ACATCCTGGGTTCTGCCCTTTGTATTTG-3’; reverse, 5’- TGACTTCCGCTTACCTTTGACTCTTTGG-3’. Identification of Cre consists of a mixture of 4 primers: primer 1, 5’-ATTTGCCTGCATTACCGGTCG-3’; primer 2, 5’-CAGCATTGCTGTCACTTGGTC-3’; primer 3, 5’-CAAATGTTGCTTGTCTGGTG-3’; primer 4, 5’-GTCAGTCGAGTGCACAGTTT-3’.

### Human samples

Written informed consents were obtained from all participants involved in this study, including 30 patients diagnosed with MDD and 30 healthy control individuals. The study was approved by the Ethics Committee of Shanghai Mental Health Center (2018-12). Clinical information, including label, gender and age was recorded for each participant, with comprehensive details available in Table S1. Blood samples were collected and promptly centrifuged at 4 °C. The resulting serum was aspirated and stored at -80 °C for subsequent analysis.

### Cell culture

*P*rimary neurons were isolated from the brains of female C57BL/6J mice at the embryonic day 16, following a previously described method (Beaudoin et al., 2012), with slight modifications. Briefly, fetal mice were subjected to the removal of the meninges, followed by the isolation of cortical and hippocampal tissues. A digestion solution was prepared using 15 mL of a mixture containing 0.125% trypsin (Gibco; #15050057) and DNase I (Sigma; #DN25) 200 U and incubated at 37 °C for 15 minutes with periodic agitation to ensure complete digestion. The digestion was terminated using FBS (Gibco; #10091-148), and the mixture was allowed to stand to sediment the cellular clumps which were then gently dissociated using a fresh culture medium, DMEM (Meilunbio; #MA0212) supplemented with 10%(v/v) FBS, with the addition of DNase I to facilitate cell dispersion. Then cells were seeded into poly-L-lysine-coated dishes or plates (Sigma; #P1399), and cultured in serum-free neurobasal medium (Gibco; #21103-049) with B27 (Gibco; #17504044), 0.5 mM glutamine (sinopharm chemical reagent; #6201083) and 1% penicillin/streptomycin (Meilunbio; #MA0110).

HEK293T cells were obtained from the Cell Bank in the Type Culture Collection Center of the Chinese Academy of Sciences and maintained in DMEM supplemented with 10% (v/v) FBS and 1% penicillin/streptomycin. All the cells were cultured in a humidified incubator at 37 °C with 5% CO_2_.

### Cannula implantation and microinjection

All stereotaxic surgeries were performed on a stereotaxic apparatus (RWD Life Science). Mice were anesthetized with Zoletil (80 mg/kg, i.p.). An incision was made on the mouse head to expose the skull surface. After scraping away the pericranium, burr holes were made on the skull. Bilateral implantation of 26-gauge guide cannulas was performed above the hippocampus (AP: −2.18 mm; ML: ±2.3 mm; DV: −2.3 mm. AP, ML and DV denote anteroposterior, mediolateral and dorsoventral distance from the bregma respectively). The cannulas were secured to the skull using stainless-steel screws and dental cement. Microinjections were made through a 33 gauge needle connected to a 10 μL Hamilton microsyringe mounted on a microinfusion pump (Harvard apparatus, USA). To prevent blockage, the needle was extended 1 mm beyond the tips of the guide cannula. For the activation or inhibition of the PI3K, 0.5 μl of 740 Y-P (30 μM in saline, administered 6 times over 2 weeks) or 0.5 μL of LY294002 (10 μM in 0.02% DMSO, administered 6 times over 3 weeks) was injected bilaterally into the hippocampus. The infusion rate of all drugs was set at 0.2 μL/min. After infusion, the needle was left in place for an additional 3 min to allow for drug diffusion. Mice were allowed to recover from anesthesia under a warming lamp to ensure their gradual return to consciousness and monitored for the first 72 h after surgery.

### Viral injection

Mice aged 8 to 12 weeks were anesthetized with Zoletil (80 mg/kg, i.p.) and positioned on the stereotactic apparatus (RWD Life Science). An incision was meticulously executed on the mouse’s scalp to reveal the underlying skull. Following the removal of the periosteum, precise trephination was conducted at selected skull sites. Adeno-associated virus (AAV, 1 μL/hemisphere) were bilaterally injected to the hippocampus with a 10 μL Hamilton syringe anchored on a microinfusion pump (Harvard apparatus, USA) at a rate of 0.2 μl/min. Viral microinjections were performed at the stereotaxic coordinate for hippocampus (AP: −2.18 mm; ML: ±2.3 mm; DV: −2.3 mm). Following infusion, the needle was left in place for additional 8 min to ensure virus diffusion and then removed carefully. After surgery, mice were allowed to recover under a warming lamp and monitored for the first 72 h after surgery.

The following AAV constructs were used: AAV2/9-CaMKIIα-ZsGreen (titre: 1.2 × 10^12^ v. g./mL, 1 μL per side bilateral into HPC, HANBIO), AAV2/9-CaMKIIα-m-Glce-3×flag-ZsGreen (titre: 1.4 × 10^12^ v. g./mL, 1 μL per side bilateral into HPC, HANBIO), AAV2/9-CaMKIIα-m-Glce-mutant-3×flag-ZsGreen (titre: 1.1 × 10^12^ v. g./mL, 1 μL per side bilateral into HPC, HANBIO), AAV2/9-CaMKIIα-EGFP-2A-3×FLAG (titre: 1.35 × 10^13^ v. g./mL, dilution: 1: 2, 1 μL per side bilateral into HPC, OBIO), AAV2/9-CaMKIIα-EGFP-P2A-NLS-Cre-WPRE (titre: 1.05 × 10^13^ v. g./mL, dilution: 1: 2, 1 μL per side bilateral into HPC, OBIO), AAV2/9-Syn-MCS-eGFP-3FLAG (1 μL per side bilateral into HPC, OBIO), AAV2/9-Syn-*Glce* shRNA-MCS-eGFP-3FLAG (1 μL per side bilateral into HPC, OBIO) and the sequence of siRNA was used as 5’-GCAAGGTGTTAGGGCTCAAAT-3’, AAV2/9-CaMKIIα-mCherry-3×FLAG-WPRE (titre: 3.99 × 10^13^ v. g./mL, dilution: 1: 10, 1 μL per side bilateral into HPC, OBIO), AAV2/9-CaMKIIα-Bdnf-3×FLAG-P2A-mCherry-WPRE (titre: 2.54 × 10^13^ v. g./mL, dilution: 1: 10, 1 μL per side bilateral into HPC, OBIO), AAV2/9-CaMKIIα- mCherry-3×FLAG-pri-mmu-mir-34c(mut 3p)-WPRE (titre: 9.25 × 10^12^ v. g./mL, dilution: 1: 2, 1 μL per side bilateral into HPC, OBIO).

### Glce knockdown and overexpression plasmids

Virus-based overexpression and knockdown of *Glce* was achieved using the pLVX-IRES-*Glce*-ZsGreen1 and PLL3.7-sh*Glce* plasmids, respectively. The sh*Glce* vector was constructed by inserting shRNA hairpin sequence into the PLL3.7 vector. We designed shRNA targeting sequences *Glce* using RNAi designer online software (http://rnaidesigner.thermofisher.com/rnaiexpress/; Invitrogen), with the sequence specified as: GCAAGGTGTTAGGGCTCAAAT. For overexpression, the full length of the mouse *Glce* sequence was amplified from cDNA using PrimeSTAR HS DNA Polymerase (Takara, R010) and inserted into pLVX-IRES-ZsGreen1 vector.

### Lentivirus-based overexpression and knockdown

Plasmids conveying either *Glce* shRNA or *Glce* overexpression sequence, along with their respective vector controls, were transfected into HEK293T cells with viral packaging vectors (psPAX2, pMD2G), using a calcium phosphate transfection method as previously described (Beaudoin et al., 2012). Before being used to affect neurons, virus was harvested from the supernatant of HEK293T cells 48 hours post-transfection. Prior to experimental use, cells were cultured in complete medium devoid of virus for at least 48 h,

### Transient transfection-mediated knockdown

The miR-34c-5p inhibitor (anti-miR-34c-5p) was procured from GenePharma Co., Ltd. (Shanghai, China). The sequence of the 2’-O-methyl-modified anti-miR-34c-5p is 5’-GCAAUCAGCUAACUACACUGCCU-3’, and the scrambled 2’-O-methyl-modified RNA sequence is 5’-CAGUACUUUUGUGUAGUACAA-3’. Transfection of primary cortical cells was performed in 12-well plates, following the protocol for Lipofectamine 2000 (Thermo Fisher, #11668019). On day in vitro 3 (DIV3), 30 pmol of RNA was mixed with 100 µl of Opti-MEM, while 1.5 µl of Lipofectamine 2000 was diluted in a separate 100 µl of Opti-MEM. After a 5-minute incubation at room temperature, the two solutions were combined and incubated for 20 minutes. The mixture was then gently added to the respective cell wells. Following a 4-hour incubation period, the medium in the wells was replaced with fresh culture medium.

### Golgi staining

Neurons were stained in vitro using the FD Rapid Golgi Stain kit, following the manufacturer’s instruction (FD Neuro Technologies, #PK401). Briefly, the mouse brain was quickly dissected and immersed into impregnation solution, followed by incubation for 14 days at room temperature in the dark. The impregnation solution was the mixture of equal volumes of solution A and B. The brains were transferred into solution C and stored at room temperature for at least 72 h in the dark, with the buffer changed at least once after the first 24 h. The brains were sectioned using a freezing microtome (Leica, Germany) at a thickness of 100 μm. Sections containing the hippocampus were mounted onto gelatin-covered microscope slides (FD Neuro Technologies, #PO101) using solution C. Sections were allowed to dry naturally for subsequent staining. They were rinsed twice in Milli-Q water for 4 min each and then placed in staining solution (solution D: solution E: Milli-Q water= 1: 1: 2). The slides were then dehydrated with gradient ethanol, vitrificated by dimethylbenzene and mounted using resinene. The stained neurons were imaged by microscope (Leica, Germany). Neurite morphology was analyzed by Image J software.

### Quantitative real-time PCR

The total RNA were extracted from tissues using FastPure Cell/Tissue Total RNA Isolation Kit V2 according to the manufacturer’s protocol (Vazyme, #RC112). Briefly, the hippocampus tissues (20 mg) or primary neurons were collected to extract total RNA. Complementary DNA (cDNA) was then generated from 500 ng of total RNA using HiScript III RT SuperMix for qPCR (Vazyme, #R323). Quantitative real-time PCRs were performed using a V7 PCR system ChamQ Universal SYBR qPCR Master Mix (Vazyme, #Q711). The Ct value was determined for the target transcripts with GAPDH used as an internal control. The relative mRNA level was calculated using the formula 2^[(Ct^ ^of^ ^gene)−(Ct^ ^of^ ^GAPDH)]^. Primer sequences used for PCR are provided in Table S2.

### Western blotting analysis

Cultured cells, mouse cortex and hippocampus tissues were lysed with RIPA buffer (0.5% Triton X-100, 0.5% deoxycholic acid sodium salt, 0.1% sodium dodecyl sulfate (SDS) and 1% phenylmethanesulfonylfluoride (PMSF) supplemented with 1% proteinase inhibitor cocktail and 1% phosphatase inhibitor) on ice for 30 min, followed by centrifugation at 12,000 × g for 10 min at 4 °C. Supernatants were separated by SDS-polyacrylamide gel electrophoresis after adjust to equal total protein content by the BCA (bicinchoninic acid) protein assay kit (Takara, #T9300A). Proteins on the gel were subsequent transferred to a polyvinyl difluoride (PVDF) membrane (PALL, #66485). After blocking with TBST buffer (20 mM Tris, pH 8.0, 150 mM NaCl and 0.1% Tween 20) containing 5% nonfat milk for 2 h, the membranes were immunoblotted with primary antibodies at 4 °C overnight, followed by probing with horseradish peroxidase (HRP)-conjugated secondary antibody. Finally, enhanced chemiluminescence (ECL) reagent was employed for signal detection. Quantification of band intensities was normalized to loading control. Data were quantified and normalized using ImageJ software and from at least three independent samples.

The antibodies used on immunoblot were purchased and diluted as follows: anti-Glce (AB-GLCERP, 1: 1000), anti-PI3K P110α (sc-293172, 1: 1000), anti-PI3K P85α (sc-1637, 1: 500), anti-PI3K P85α (13666, 1: 1000), anti-PI3K P85α (ab191606, 1: 1000), anti-BDNF (ab108319, 1: 1000), anti-Akt (9272, 1: 1000), anti-Phospho-Akt (4060, 1: 1000), anti-TrKB (4603, 1: 1000), anti-Phospho- TrKB (ABN1381, 1: 500), HRP- conjugated Affinipure Goat Anti-Mouse (SA00001-1, 1: 5000), HRP-conjugated Affinipure Goat Anti-Rabbit (SA00001-2, 1: 5000), anti-GAPDH (60004-1-Ig, 1: 5000).

### Immunofluorescence

Mice that underwent behavioral analysis were deeply anesthetized with Zoletil and perfused transcardially with saline, followed by 4% PFA in 0.1 M PBS. Brains were then removed, post-fixed overnight in 4% PFA at 4 °C, and transferred into 30% sucrose in 0.1 M PBS for dehydration. Mouse brain sections containing hippocampus were cut into 30 μm thick and stored in 0.1 M PBS. Then brain sections or the primary neurons were fixed in 4% paraformaldehyde (PFA)/PBS for 15 min. Samples were washed twice with PBS, permeabilized for 20 min with PBS/0.3% Triton X-100 (PBST), blocked for at least 1 h at room temperature (RT) with 5 % bovine serum albumin (BSA) in PBST, and incubated in primary antibodies overnight at 4 °C. After washing with PBS, the samples were incubated with fluorescence conjugated secondary antibody. All the slices were counterstaining with DAPI. After mounting, images were taken using confocal laser scanning microscope (Lecia, Lecia TCS SPS CFSMP, Germany) and Leica Application Suite X software. Measurements of neurite were analyzed using Image-Pro Plus software.

The antibodies used on immunofluorescence were purchased and diluted as follows: anti-Glce (PA559194, 1: 200), anti-PI3K P110a (PA5-18610, 1: 100), anti-PI3K P85a (AF2998, 1: 100), anti- PI3K P110a (sc-293172, 1: 100), anti-PI3K P85a (sc-1637, 1: 50), anti-GM130 (ab169276, 1: 200), anti-BDNF (ab108319, 1: 500), anti-NeuN (MAB377, 1: 500), anti-MAP2 (AB5622, 1: 500), anti-CaMKIIa (66843-1-lg, 1: 200), anti-Iba1 (ab178846, 1:500), anti-GAD67 (ABclonal, 1: 100), Goat anti-Rabbit IgG Alexa Fluor™ 594 (A11037, 1: 200), Goat anti-Rabbit IgG Alexa Fluor™ 488 (Cat# A-11008, 1: 200), Goat anti-Mouse IgG Alexa Fluor™ 488 (A-11029, 1: 200), Goat anti-Rabbit IgG Alexa Fluor™ Plus 594 (A32740, 1: 200), Donkey anti-Goat IgG Alexa Fluor™ 594 (A-11058, 1: 200), Goat anti-Rabbit IgG Alexa Fluor™ 647 (A-21246, 1: 200).

### PI3K enzymatic activity test

The PI3K enzymatic activity test was conducted according to the manufacturer’s instruction (GENMED, #GMS50058.2). For each experimental group, 10 mg hippocampal tissue was excised and homogenized in solution B at 4 °C. The lysate was subjected to vigorous shaking for three intervals, each lasting 30 seconds in duration, followed by centrifugation at 10000 g for 10 minutes at 4 °C. The supernatant was collected, and protein concentrations were measured and adjusted to achieve equilibration. The prepared samples were then transferred into sterile 1.5 mL tubes for subsequent PI3K enzymatic activity test. In a 96-well plate, the reaction mixture was prepared by adding 65 μL of solution C, 10 μL of solution D, 10 μL of solution E, and 10 μL of solution F, and incubated at 30 °C for 30 minutes. Subsequently, the wells were supplemented with either the test samples or solution G as a negative control. Immediately, absorbance readings at 340 nm were taken at 0 and 5 minutes post-incubation. The PI3K enzymatic activity was defined as [(Sample Absorbance−Background Absorbance) × 0.1 (Assay Volume, mL) × Sample Dilution Factor] / [(Optical path length × 6.22 (Molar Absorptivity Coefficient) × 5 (Reaction Time, min))] / (protein concentration of samples, mg/mL) = μmol NADH/min/mg.

### Co-immunoprecipitation

Co-immunoprecipitation (Co-IP) was performed using primary cortical neurons and hippocampal tissues from C57BL6 mice. Samples were lysed on ice for 30 min using IP lysis buffer containing 1% phenylmethanesulfonylfluoride (PMSF) supplemented with 1% proteinase inhibitor cocktail, and 1% phosphatase inhibitor. After centrifuged at 12,000 g for 10 min at 4 °C, supernatants were incubated overnight at 4 °C with anti-*Glce* antibody (AB-GLCERP, 1: 100), anti-PI3K p110α antibody (sc-293172, 1: 50) or anti-PI3K p85α antibody (sc-1637, 1: 50). Protein A/G agarose (30 μL each) were added to capture the antibody-protein complexes, followed by an overnight incubation at 4 °C. Samples precipitated with nonspecific IgG (3900, 1: 250) were used as negative control. The beads with protein complexes were washed and collected. The immunoprecipitated complex were boiled with 1 × SDS buffer at 95 °C and loaded to 10% SDS-PAGE gel, followed by the immunoblotting. For detailed procedural instructions, refer to the manual in the COIP kit (Absin, #abs955).

### Surface plasmon resonance (SPR) analysis

SPR experiment was performed using a BIACORE T200 system (GE Healthcare, Stockholm, Sweden). PI3K proteins (p110 alpha/p85 alpha) were immobilized on a CM5 sensor chip by the amine coupling method. For interaction measurements, different concentration of *Glce* protein was injected into the chips. All procedures were conducted in HBS-EP (0.15 M NaCl, 0.01 M HEPES, 3 mM EDTA and 0.005% surfactant P20, pH 7.4) running buffer. Kinetic parameters were analyzed using 1: 1 binding model by BIACORE T200 Evaluation Software Version 1.0.

#### Protein-protein docking

The crystal structure of dimeric human *Glce* (37) (PDB Code: 6HZZ) and the AlphaFold3 (Abramson et al., 2024) -predicted model of hPI3K, comprising the subunits p110a and p85a, were preprocessed utilizing the Protein Preparation Wizard (Sastry et al., 2013) implemented in the Schrödinger suite. The co-crystallized ligands included in the structure were deleted and hydrogen atoms as well as the missing side chains of residues were added. Structure refinement was performed using the OPLS3 force field (Harder et al., 2016) with harmonic restraints applied to heavy atoms. The optimized structures of dimeric h*Glce* and hPI3K were subsequently submitted to the ClusPro Server (Kozakov et al., 2017) for protein-protein docking, generating 88 complex models. These models were evaluated using Rosetta (Alford et al., 2017) energy scoring functions. The optimal binding pose was selected based on both energetic considerations and conformational plausibility.

### Dual-luciferase reporter gene assay

The gene fragment containing the *Glce* 3’ untranslated region (UTR) (3’UTR-*Glce*) and 3’UTR-*Glce* mutant was cloned into the pmiR-RB-ReportTM vector (Ribobio Co., Ltd., Guangzhou, China). HEK293T cells were seeded in a 96-well plate at a density of 5 × 10^4 cells per well and cultured overnight to achieve 70% confluence the following day. According to the manufacturer’s instructions for Lipofectamine^TM^ 2000, miR-34c-5p or anti-miR-34c-5p, along with either pmiR-RB-3’UTR-*Glc*e, pmiR-RB-3’UTR-*Glce* mutant or pmiR-RB-3’UTR-NC, were co-transfected into the cells and incubated for an additional 36 hours. The dual-luciferase reporter assay was performed using the Dual Luciferase Assay System kit (Promega, #E1910), following the procedural steps outlined in the kit’s manual.

### Behavioral procedures

#### Chronic restrained stress (CRS) paradigm

Male C57BL6/J mice, aged 8 to 12 weeks, were randomly divided into 2 groups. Mice in the experiment group were restrained in a 50-mL conical tube with hole for airflow for 3 hours per day lasting for 3 weeks, and mice in the control group were deprived of food and water at the same time as the experiment group (Yang et al., 2018). Brain tissues were dissected in 24 hours after the final behavior test.

#### Chronic social defeat stress (CSDS) paradigm

Chronic social defeat stress (CSDS) paradigm was conducted according to previously described methods, with slight modifications (Golden et al., 2011). The CSDS protocol involved the following steps: firstly, CD-1 aggressor mice were screened for three consecutive days with strict criteria: CD-1 mice were required to attack the C57BL/6 mouse for at least five consecutive sessions within 3 min, and the latency to initial aggression had to be less than 60 s. Following the screening, CD-1 mice were individually housed in cages separated into two compartments by perforated plastic separators. The C57 mice were placed into the homecage of CD-1 mice and physically attacked by aggressive CD-1 for 10 min. The mice were separated by a perforated plexiglass partition and maintained in sensory contact for another 24 h after 10 min of physical contact. After 10 days of CSDS, the mice were housed individually. Control animals were housed in pairs, each on either side of the perforated plexiglass partition, and were never exposed to physical or sensory contact with CD-1 mice before the social interaction test. 24 h after the last social defeat stress, behavioral assessments were applied to evaluate depressive-like behaviors in mice. Brain tissues were dissected within 24 hours after the final behavioral test.

#### Social interaction test (SIT)

Social interaction behavior was assessed using a two-stage social interaction test. In the initial 3 min test (no target), the experimental C57 mouse explored a square shaped arena (40 × 40 × 40 cm^3^) containing an empty wire mesh cage (10 × 6.5 × 20 cm^3^) placed on one side. In the subsequent 3 min test (target), the experimental C57 mouse was reintroduced into the arena with an unfamiliar CD-1 mouse in the wire mesh cage. The time of the experimental mouse spent in the “Interaction Zone” (a rectangular area (14 × 24 cm^2^) around the enclosure that was applied to display the target CD-1 mice) and “Corner Zone” (9 × 9 cm^2^ zone projecting from both corner joints opposing the enclosure) were quantified by the video tracking software. The social interaction index was calculated as the time spent in interaction zone with the target divided by the time spent in interaction zone without the target.

#### Tail suspension test (TST)

The tail suspension test was performed according to previous protocol (Castagné et al., 2010). Briefly, mice were suspended by their tails approximately 70 cm above the ground using adhesive tape applied 1 cm from the tail tip. Their behavior was video-recorded for 6 minutes. Immobility time, defined as remaining motionless, was measured during the last 4 minutes. Investigators were blinded to the treatment allocation during behavior assessment.

#### Forced swimming test (FST)

Individual mice were placed in a cylinder (25 cm height; 15 cm diameter) filled with water (23–25 °C) and allowed to swim for 6 min (Castagné et al., 2010). The depth of water (15 cm height) prevented animals from touching the bottom with their tails or hind limbs. Video recordings captured animal behavior from the side. Immobility time during the last 4 minutes was quantified offline by an observer blinded to treatment allocation. Immobility was defined as remaining floating or motionless, with only movements necessary for balance.

#### Sucrose preference test (SPT)

Mice underwent habituation to two bottles of drinking water for three days, during this period, mice position preference was ascertained by monitoring and recording the water consumption from two bottles. Mice then were habituated with drinking water and 1.0% sucrose solution for 12 h and reversing the bottle position for habituation for 12 h. After habituation, mice were deprived of water for 24 h and then offered a choice between drinking water and 1.0% sucrose solution in the dark phase for 2 h, with the sucrose solution placed on the non-preferred side of the mouse’s enclosure. Total consumption of each fluid was measured, and the sucrose preference score was calculated as: amount of sucrose consumed × 100% / (sucrose consumed + water consumed).

#### Locomotor activity test (LAT)

Mice were assessed for locomotor activity over 10 min in an open field arena (40 × 40 × 40 cm^3^). Motion tracks of the mice were recorded by the video camera, and analyzed using Shanghai Jiliang Software. Locomotor activity was evaluated as the total distance traveled.

#### Quantification and statistical analysis

All data were presented as means ± standard error of the mean (S.E.M.). The number of experimental replicates (n) was specified in the figure legends, indicating the number of experimental subjects independently treated in each condition. Statistical comparisons were analyzed using GraphPad Prism (version 10.0), employing appropriate methods as described in the figure legends. Single-variable differences were assessed using two-tailed unpaired Student’s *t* tests, while grouped differences were analyzed with One- or Two-way analysis of variance (ANOVA) followed by Bonferroni post-hoc test. Statistical significance was set at *P < 0.05, **P < 0.01, ***P < 0.001, ****P < 0.0001.

## Supporting information

Graphical Abstract

## Data availability

All data reported in this paper will be shared by the lead contact upon request. Any additional information required to reanalyze the data reported in this paper is available from the lead contact upon request.

## Code availability

This paper does not report original code or mathematical algorithm.

## Author contributions

KD, JGL, YJW, MC, LH, SY and JL conceived the project, conceptualized the experiments and wrote the manuscript, which was assisted by XL, YX, and YL (Yang Li). MC performed most of the experiments and analyzed all the data assisted by LH, SY, JL, FH, YZ (Yiwen Zhang), ZD and WL. YL (Yanling Li), FG, TL, JD and YZ (Yu Zhang) assisted with animal behavior experiments and provided technical help. MX assisted the protein-protein docking experiment and wrote this part of the manuscript. XL, and JN provided the serum of human. JF performed the construction of transgenic mice. CJ drew the schematic illustration.

## Acknowledgments

We thank Dr. Haiyan Zhang (Shanghai Institute of Materia Medica, Chinese Academy of Sciences, Shanghai, China) and Dr. Huan Wang (Shanghai Institute of Materia Medica, Chinese Academy of Sciences, Shanghai, China) for guiding methods of neuron culture.

---

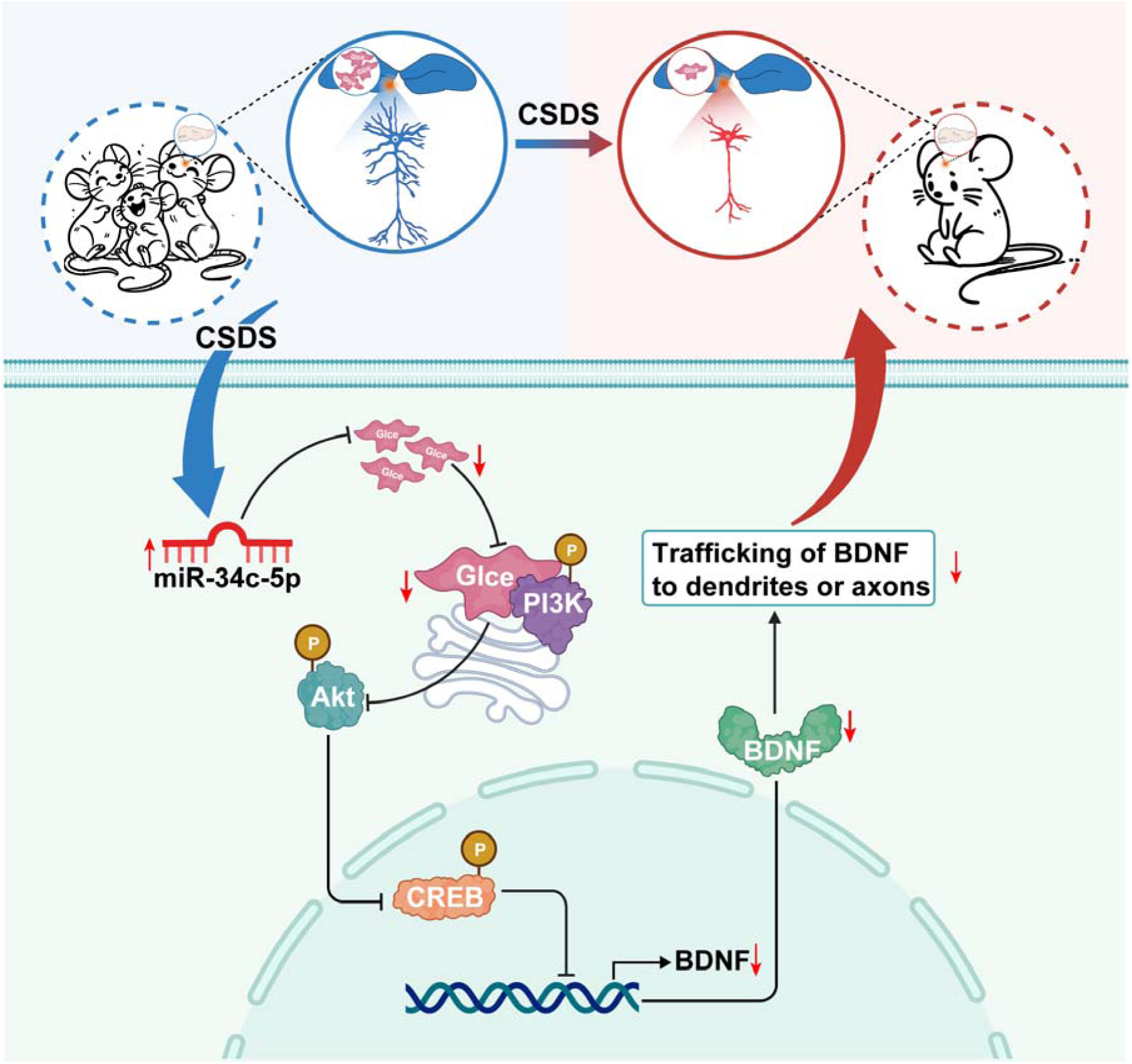

