## Supplementary materials for "Hippocampal Glucuronyl C5-epimerase promotes stress resilience by directly engaging PI3K through a non-enzymatic mechanism"

*Chen et al.*

**Table of Contents**

1. **Supplemental Tables** (Page 3-4)

Supplemental Table 1. Parameters of human subjects for serum *Glce* level detection.

Supplemental Table 2. PCR primers used in this study.

2. **Supplemental Figures** (Page 5-13)

Supplemental Figure 1. *Glce* is highly expressed in hippocampal excitatory glutamatergic neurons.

Supplemental Figure 2. *Glce* is significantly reduced in NKO and hNKO mice.

Supplemental Figure 3. Knockdown of *Glce* in the hippocampus does not induce depressive-like behaviors in mice.

Supplemental Figure 4. Overexpression of either *Glce* or BDNF can rescue neuronal atrophy induced by *Glce* loss.

Supplemental Figure 5. Deletion of *Glce* in the hippocampus decreases BDNF expression.

Supplemental Figure 6. *Glce* interacts with PI3K and regulates the PI3K-AKT-CREB pathway.

Supplemental Figure 7. *Glce* expression is downregulated in the hippocampus of mice with CRS-induced depression.

Supplemental Figure 8. *Glce* is significantly decreased in male patients with MDD.

Supplemental Figure 9. Overexpression of *Glce* and mut*Glce* maintains BDNF expression under CSDS stimulation.

| 1. **Supplemental Tables**   **Supplemental Table 1. Parameters of human subjects for serum *Glce* level detection.** | | | | | | | |
| --- | --- | --- | --- | --- | --- | --- | --- |
| **Healthy** |  | |  |  | **MDD** |  | |
| **Sample ID** | **Gender** | **Age** | |  | **Sample ID** | **Gender** | **Age** |
| A3108S | F | 80 | |  | GZ001 | F | 66 |
| A3130S | F | 72 | |  | GZ002 | F | 71 |
| A7319S | F | 74 | |  | GZ003 | M | 62 |
| A7326S | F | 73 | |  | GZ004 | M | 76 |
| A7331S | M | 71 | |  | GZ005 | F | 67 |
| A7354S | F | 74 | |  | GZ006 | F | 73 |
| A7406S | F | 73 | |  | GZ007 | F | 71 |
| A7407S | M | 72 | |  | GZ008 | F | 60 |
| A7416S | M | 74 | |  | GZ009 | M | 70 |
| A7508S | F | 72 | |  | GZ010 | M | 73 |
| A7517S | M | 71 | |  | GZ011 | F | 81 |
| A7617S | F | 72 | |  | GZ013 | M | 74 |
| A7637S | M | 72 | |  | GZ014 | F | 75 |
| A7644S | F | 71 | |  | GZ015 | F | 69 |
| A7654S | F | 74 | |  | GZ018 | M | 64 |
| A7657S | F | 73 | |  | GZ019 | F | 75 |
| A7688S | M | 71 | |  | GZ020 | F | 73 |
| A7707S | F | 73 | |  | GZ021 | F | 74 |
| A7708S | F | 69 | |  | GZ022 | F | 77 |
| A7724S | F | 74 | |  | GZ023 | F | 59 |
| A7742S | F | 73 | |  | GZ025 | F | 70 |
| A7747S | M | 73 | |  | GZ026 | F | 77 |
| A7761S | M | 73 | |  | GZ028 | F | 66 |
| A7763S | F | 66 | |  | GZ029 | F | 66 |
| A7366S | F | 72 | |  | GZ030 | F | 65 |
| A7436S | F | 73 | |  | GZ031 | F | 72 |
| A7723S | F | 74 | |  | GZ032 | M | 74 |
| A7687S | F | 71 | |  | GZ033 | M | 62 |
| A7451S | M | 73 | |  | GZ034 | F | 85 |
| A7501S | F | 72 | |  | GZ012 | F | 64 |

| **Supplemental Table 2. PCR primers used in this study.** | | |
| --- | --- | --- |
| Name | Sequence | Supplier |
| *GAPDH*-Forward Primer | AGTGCCAGCCTCGTCCCGTAG | Sangon |
| *GAPDH*-Reverse Primer | GTGCCGTTGAATTTGCCGTGAGTG | Sangon |
| *BDNF*-Forward Primer | AGGTCTGACGACGACATCACT | Sangon |
| *BDNF*-Reverse Primer | CTTCGTTGGGCCGAACCTT | Sangon |
| *Glce*-Forward Primer | GCAGCTCGGGTCAACTATAAG | Sangon |
| *Glce*-Reverse Primer | CTGAATCCACTACTCAAGTGCC | Sangon |
| *U6*-Forward Primer | CTCGCTTCGGCAGCACA | Sangon |
| *U6*-Reverse Primer | AACGCTTCACGAATTTGCGT | Sangon |
| *miR-34c-5p*-Forward Primer | CGCGAGGCAGTGTAGTTAGCT | Sangon |
| *miR-34c-5p*-Reverse Primer | AGTGCAGGGTCCGAGGTATT | Sangon |

2. **Supplemental Figures**

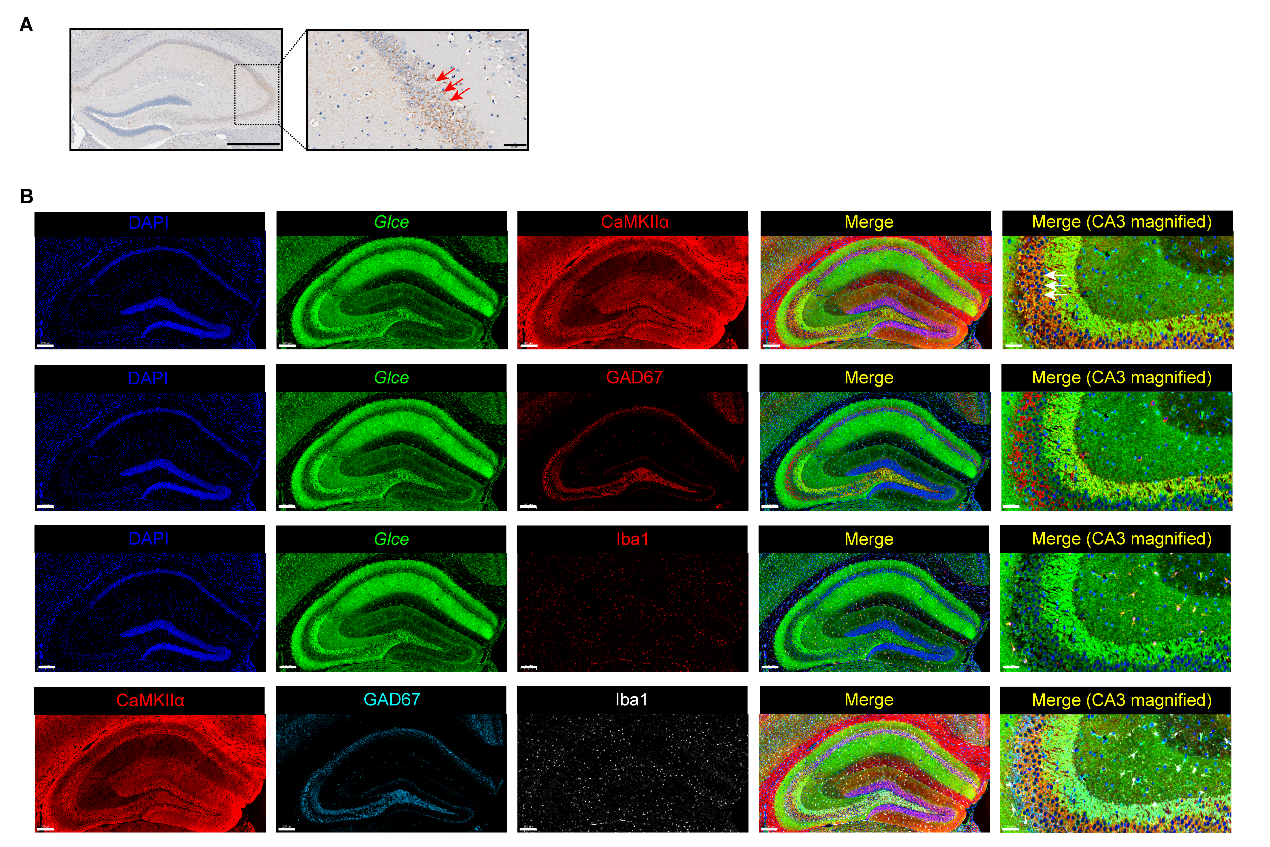

**Supplemental Figure 1. *Glce* is highly expressed in hippocampal excitatory glutamatergic neurons.** (**A**) Immunochemistry images showing *Glce* expression in the hippocampus of C57BL/6 mice. Scale bars, 500 μm, magnified image, 50 μm. (**B**) Immunofluorescence showing *Glce* (green) expression in the excitatory glutamatergic neurons (marked by CaMKIIα), GABAergic neurons (labeled by GAD67), Microglia cells (represented by Iba1) of hippocampus of C57BL/6 mice. Note: In the five-color co-staining figure, panels DAPI, *Glce*, CaMKIIα, GAD67 and Iba1 are both from the same stained slice, with different colors applied only for illustrative purposes. Scale bars, 200 μm; CA3 magnified image 50 μm.

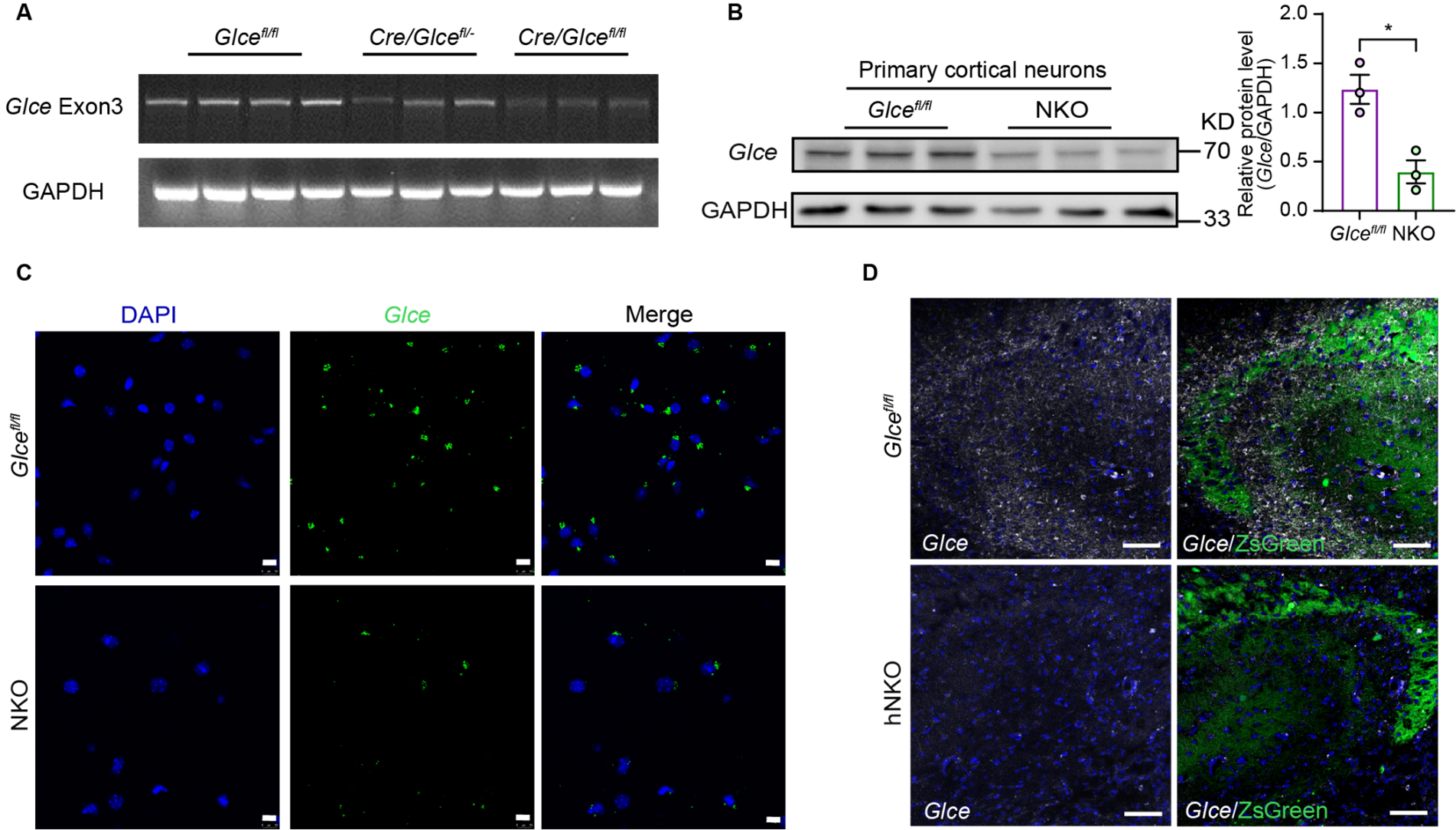

**Supplemental Figure 2. *Glce* is significantly reduced in NKO and hNKO mice.** (**A**) Verification of knockout sequence and Cre enzyme activity of *Glce* NKO (Cre/*Glce^fl/fl^*) mice by PCR. GADPH was used as internal control. (**B**) *Glce* protein expression was detected in primary cortical neurons via western blot (Left). Quantification of *Glce* protein expression levels (Right). n = 3 for *Glce^fl/fl^* mice and n = 3 for NKO mice. (**C**) Immunofluorescence showing *Glce* (green) expression in the primary hippocampal neurons cultured for 3 days of *Glce^fl/fl^* and NKO mice. Scale bars, 10 μm. (**D**) Immunofluorescence showing *Glce* (white) expression in the hippocampus of *Glce^fl/fl^* mice and hNKO mice with ZsGreen fluorescence marking hippocampal pyramidal neurons. Scale bars, 50 μm. Two-tailed unpaired *t* test. Data were presented as mean ± SEM. *P < 0.05.

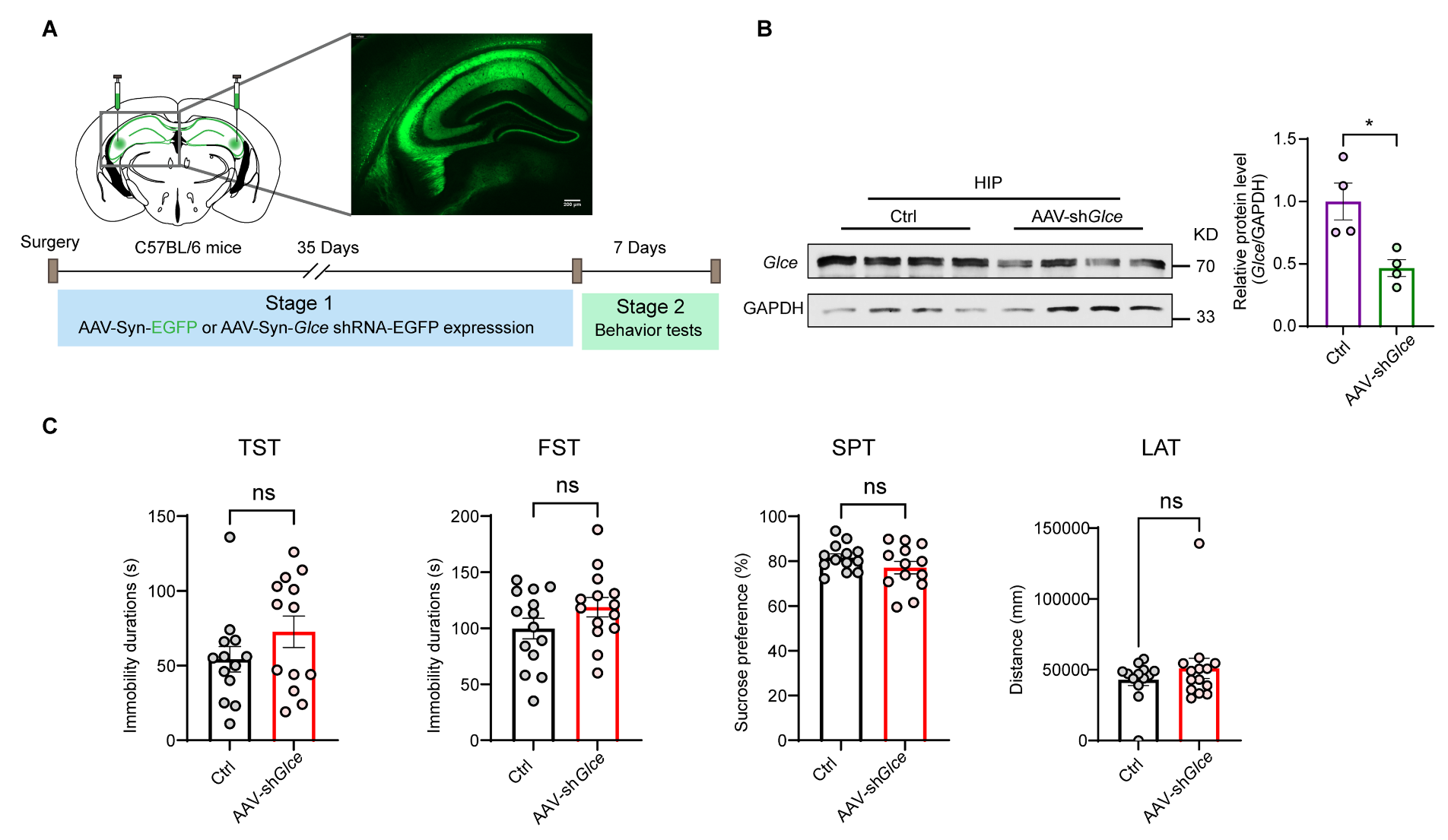

**Supplemental Figure 3. Knockdown of *Glce* in the hippocampus** **does not induce depressive-like behaviors in mice.** (**A**) Schematic construction of postnatally neuronal *Glce* knockdown in the HPC via bilateral brain stereotactic injection of AAV-syn-*Glce* shRNA- EGFP and AAV-syn-EGFP as Ctrl in 8- to 12-week-old male C57BL/6 mice. After 35 days virus expression, depression-related behavior tests were conducted. Representative image showing expression of EGFP fluorescence in the HPC (top). Scale bars, 200 μm. (**B**) Western blot (left) and quantification (right) showing *Glce* protein expression in the hippocampus from sh*Glce* mice. n = 4 for Ctrl mice and n = 4 for sh*Glce* mice. Two-tailed unpaired *t* test. Data were represented as mean ± SEM; *P < 0.05. (**C**) Effect of hippocampal *Glce* knocked down on depression-related behaviors as evaluated by TST; FST; SPT and LAT. n = 13, 13 in the TST; n = 14, 14 in the FST; n = 13, 13 in the SPT and n = 13, 14 in the LAT for Ctrl and sh*Glce* group, respectively. Two-tailed unpaired *t* test or Mann-Whitney test. Data were represented as mean ± SEM; ns, not significant, *P < 0.05.

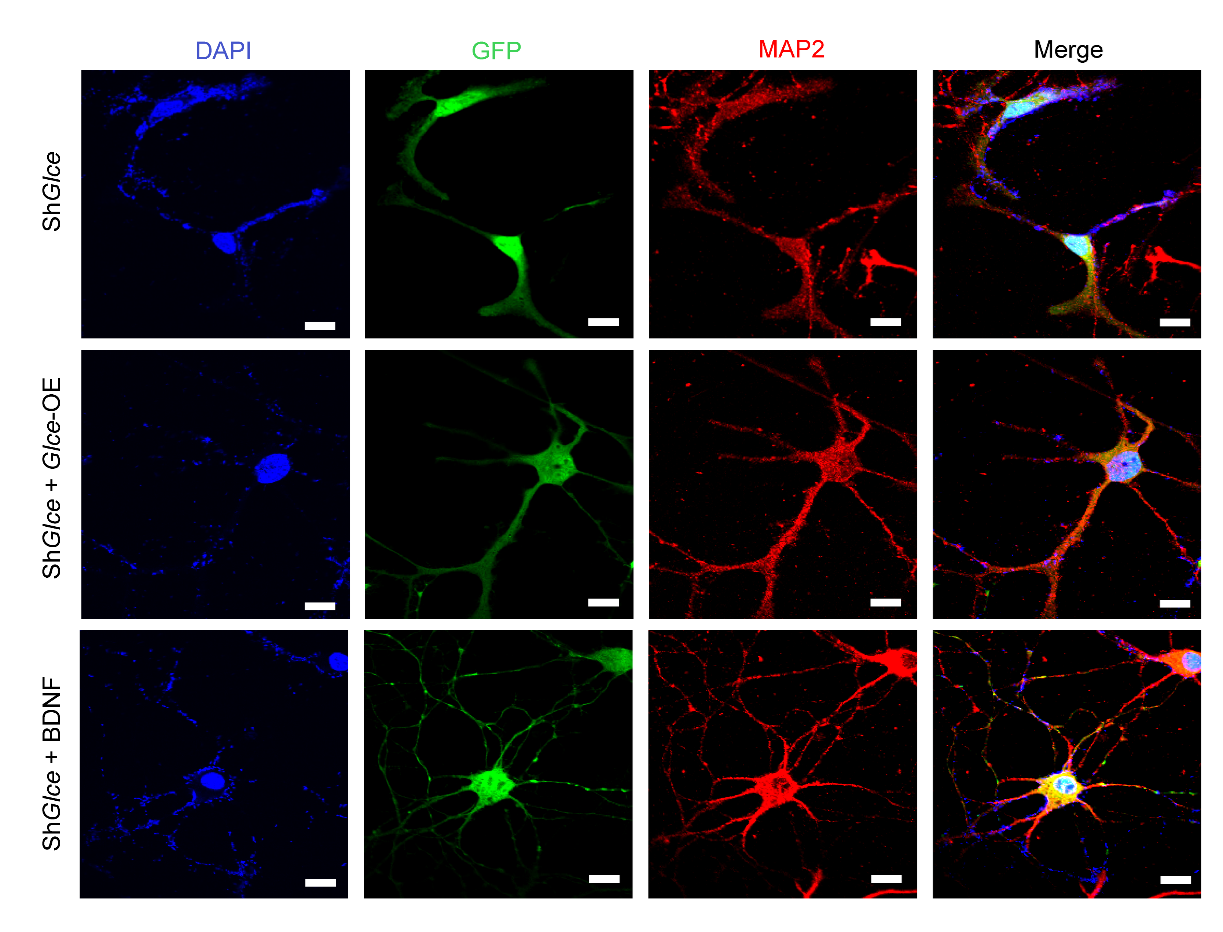

**Supplemental Figure 4. Overexpression of either *Glce* or BDNF can rescue neuronal atrophy induced by *Glce* loss.** Primary hippocampal neurons derived from C57BL/6J mice were transfected with *Glce* shRNA (PLL3.7-sh*Glce*) at DIV3. Regarding Sh*Glce + Glce*-OE group, *Glce* shRNA and *Glce*-OE lentiviruses were co-transfected into the neurons. For Sh*Glce +* BDNF group, recombinant human BDNF protein (20 ng/mL) was added into the cells transfected with sh*Glce* at DIV5. After a 7-day culture period, treated neurons were used for immunofluorescence. MAP2 antibody was employed to depict neuronal morphology. Scale bars, 100 μm.

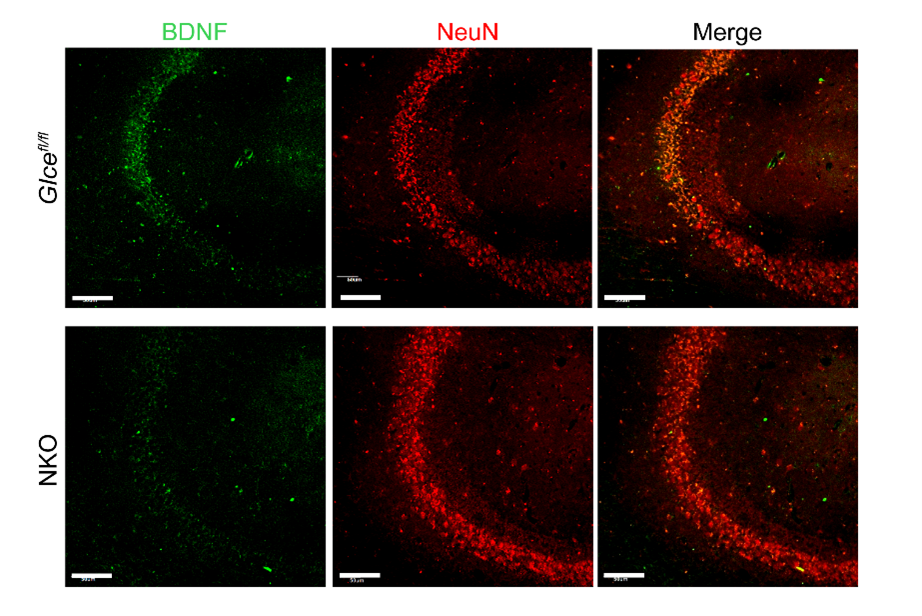

**Supplemental Figure 5. Deletion of *Glce* in the hippocampus decreases BDNF expression.** Immunofluorescence showing the BDNF (green) expression in the hippocampal CA3 region in *Glce* NKO mice and littermate *Glce^fl/fl^* mice. Neuronal nuclei (NeuN, red) was employed to specifically mark mature neurons. Scale bars, 100 μm.

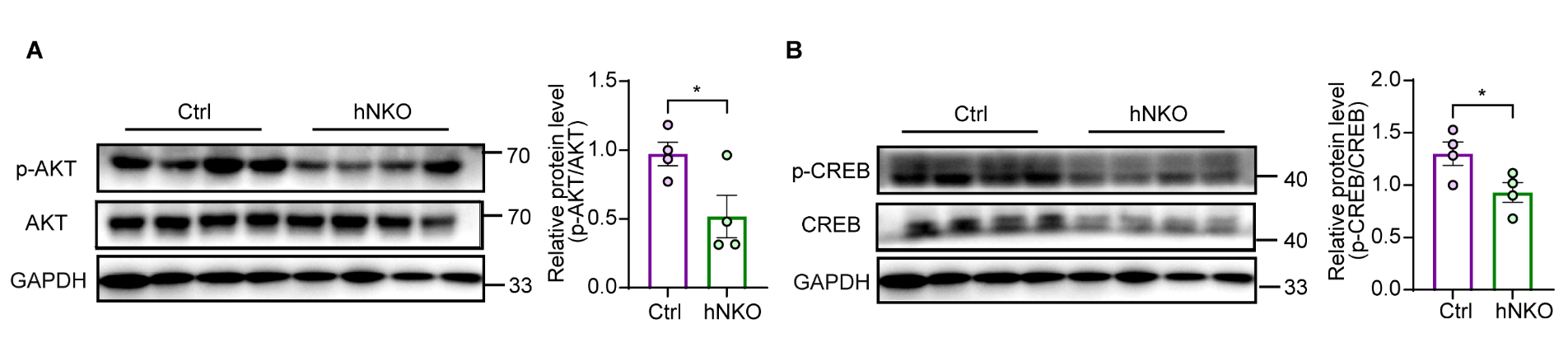

**Supplemental Figure 6. *Glce* interacts with PI3K and regulates the PI3K-AKT-CREB pathway.** (**A**) Western blot (left) and quantification (right) results indicating p-AKT/AKT expression levels in hNKO mice. GAPDH was used as an internal reference control. Two-tailed unpaired *t* test. Data were presented as mean ± SEM. *P < 0.05. (**B**) Western blot (left) and quantification (right) results showing p-CREB/CREB expression levels in hNKO mice. GAPDH was used as an internal reference control. Two-tailed unpaired *t* test. Data were presented as mean ± SEM. *P < 0.05. Note: Figure 3D, Supplemental Figure 6, A and B were derived from the same Western blot experiment with common internal control GAPDH band.

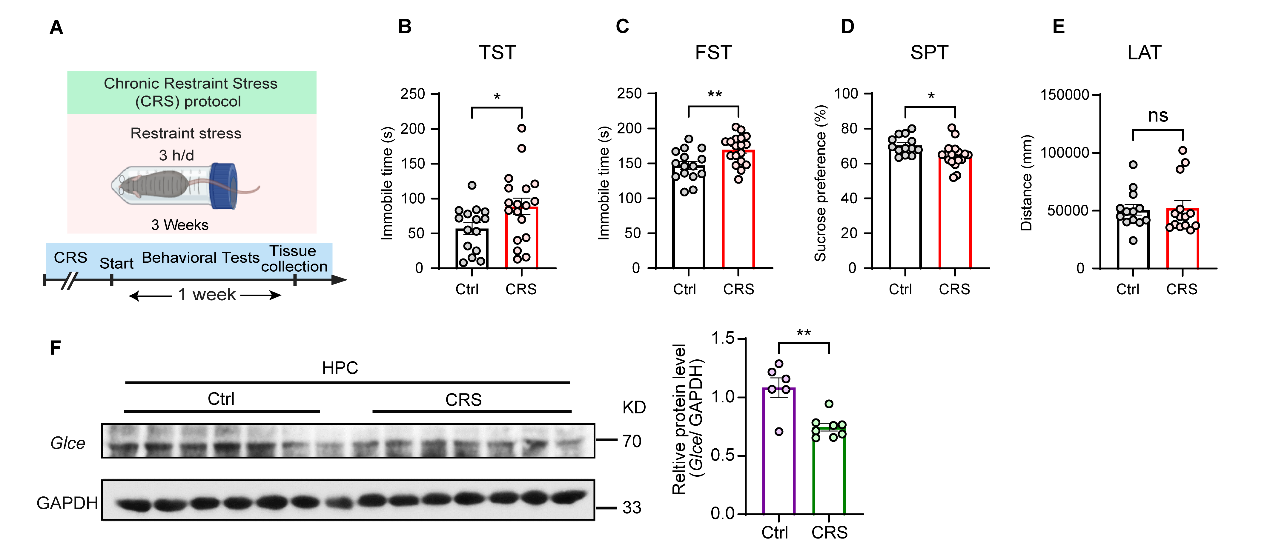

**Supplemental Figure 7. *Glce* expression is downregulated in the hippocampus of mice with CRS-induced depression.** (**A**) Schematic of the CRS paradigm revealing that male C57BL/6 mice were restrained in a 50-mL conical tube with hole for airflow for 3 hours per day lasting for 3 weeks. Depressive-related behavior tests were examined to validate the success of CRS model. (**B** to **E**) Depression-related behavior tests performed in CRS mice by TST (**B**); FST (**C**); SPT (**D**) and LAT (**E**). n = 15, 18 mice in the TST; n = 15, 18 mice in the FST; n = 13, 18 mice in the SPT and n = 13, 14 mice in the LAT for Ctrl and CRS group. Two-tailed unpaired *t* test, followed by Mann-Whitney test. Data were represented as mean ± SEM. ns, not significant, *P < 0.05, **P < 0.01. (**F**) Western blot (left) and quantification (right) showing *Glce* protein expression in the hippocampus (HPC) from CRS model mice. GAPDH was used as an internal reference control. n = 6 for Ctrl mice and n = 8 for CRS-treated mice. Two-tailed unpaired *t* test or Mann-Whitney test. Data were presented as mean ± SEM. **P < 0.01.

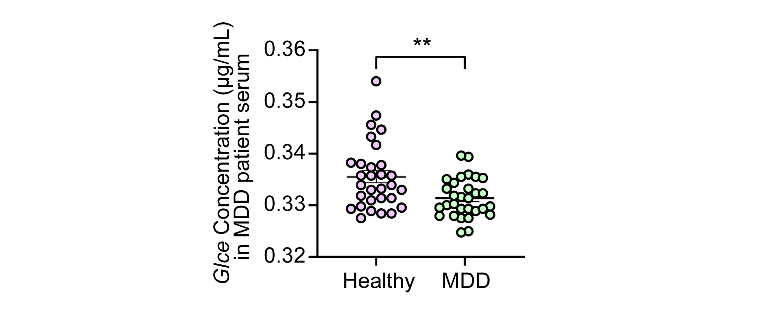

**Supplemental Figure 8. *Glce* is significantly decreased in male patients with MDD.** ELISA assay showing plasma *Glce* protein levels in MDD patients and relative healthy individuals, n = 30, 30 humans. Two-tailed unpaired *t* test or Mann-Whitney test. Data were presented as mean ± SEM. **P < 0.01.

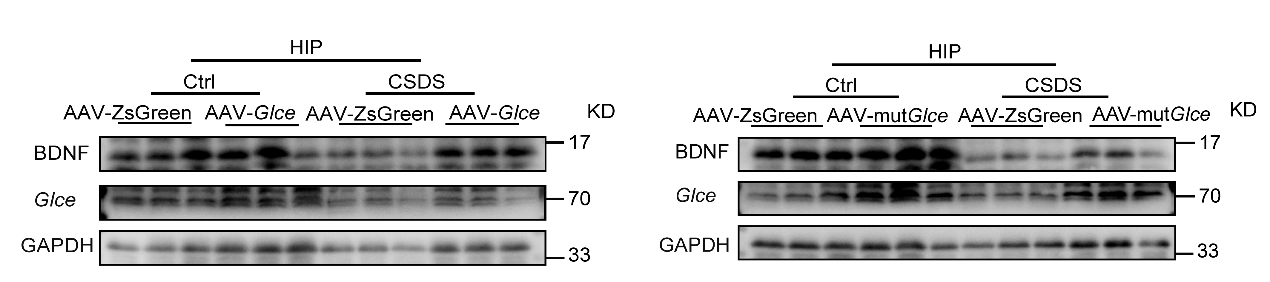

**Supplemental Figure 9. Overexpression of *Glce* and mut*Glce* maintains BDNF expression under CSDS stimulation.** Effect of *Glce* and *Glce* mutant overexpression on BDNF expression in CSDS stimulated C57BL/6J mice. n = 3, 3, 3, 3 in Ctrl + AAV-ZsGreen, Ctrl + AAV-*Glce*, CSDS + AAV-ZsGreen and CSDS + AAV-*Glce* group (left); n = 3, 3, 3, 3 in Ctrl + AAV-ZsGreen, Ctrl + AAV-mut*Glce*, CSDS + AAV-ZsGreen and CSDS + AAV-mut*Glce* group (right).
