## Supplementary figures and images for "Hippocampal Glucuronyl C5-epimerase promotes stress resilience by directly engaging PI3K through a non-enzymatic mechanism"

### Graphical Abstract

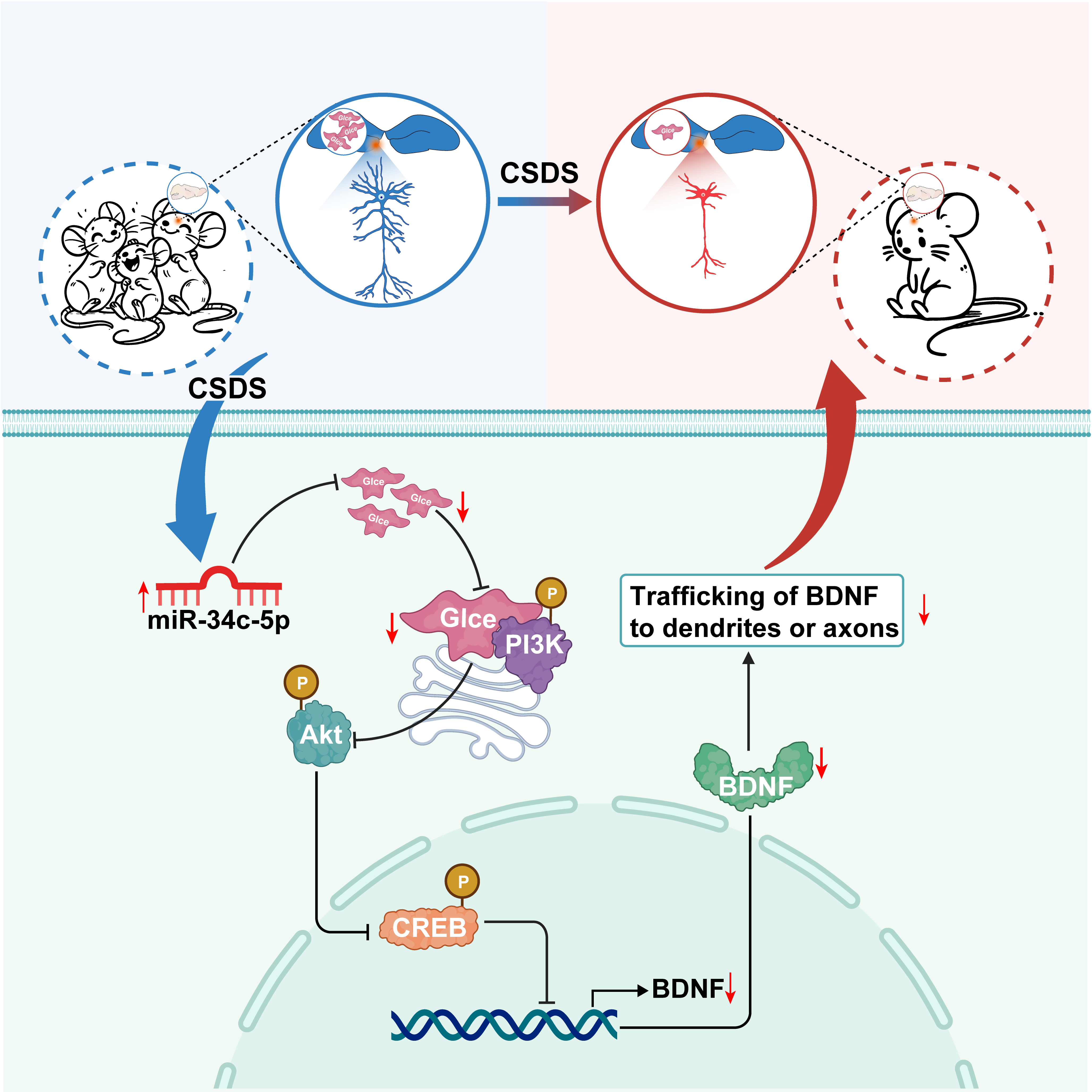
